# MGIDI: towards an effective multivariate selection in biological experiments

**DOI:** 10.1101/2020.07.23.217778

**Authors:** Tiago Olivoto, Maicon Nardino

## Abstract

Multivariate data are common in biological experiments and using the information on multiple traits is crucial to make better decisions for treatment recommendations or genotype selection. However, identifying genotypes/treatments that combine high performance across many traits has been a challenger task. Classical linear multi-trait selection indexes are available, but the presence of multicollinearity and the arbitrary choosing of weighting coefficients may erode the genetic gains. We propose a novel approach for genotype selection and treatment recommendation based on multiple traits that overcome the fragility of classical linear indexes. Here, we use the distance between the genotypes/treatment with an ideotype defined *a priori* as a multi-trait genotype-ideotype distance index (MGIDI) to provide a selection process that is unique, easy-to-interpret, free from weighting coefficients and multicollinearity issues. The performance of the MGIDI index is assessed through a Monte Carlo simulation study where the percentage of success in selecting traits with desired gains is compared with classical and modern indexes under different scenarios. Two real plant datasets are used to illustrate the application of the index from breeders and agronomists’ points of view. Our experimental results indicate that MGIDI can effectively select superior treatments/genotypes based on multi-trait data, outperforming state-of-the-art methods, and helping practitioners to make better strategic decisions towards an effective multivariate selection in biological experiments.

## 1. Introduction

Experienced breeders often keep in mind a set of plant traits that, if brought together in one new genotype would lead to high performance. Here, this target genotype is treated as *ideotype*, concept introduced by Donald (1968) nearly a half-century ago. For wheat, for example, breeders often look for healthily- and high-yielding plants, that provide grains with high baking quality. The idea behind ideotype-design is increasing crop performance, of course, focusing on the selection of genotypes based on multiple traits simultaneously.

Several linear selection indexes (Cerón-Rojas and Crossa, 2018) help breeders to select superior genotypes. A widely-used base linear phenotypic selection index (Bhering et al., 2012; Bizari et al., 2017; Jahufer and Casler, 2015; Burdon and Li, 2019) is the Smith-Hazel (SH) index (Smith, 1936; Hazel, 1943). To compute the SH index, breeders use the phenotypic and genotypic (co)variance matrices as well as a vector of economic weights to determine how a vector of index coefficients has to be chosen to maximize the correlation of unknown genetic values and phenotypic values. Because the SH index requires inverting a phenotypic covariance matrix (Smith, 1936), the presence of multicollinearity –which will certainly appear when several traits are assessed– can result in poorly conditioned matrices and biased index coefficients, thus, affecting the estimates of genetic gains. Besides the multicollinearity issue, breeders often face difficult choices to express the economic value of traits, converting them into realistic economic weightings (Bizari et al., 2017).

Even though the SH index is widely used as a multi-trait selection index, there has been evidence that the adoption of this index is not interesting for plant breeding, either in early plant breeding trials (Bhering et al., 2012) or in advanced stages of breeding programs (Jahufer and Casler, 2015), which are often conducted in multi-environment trials (Olivoto et al., 2019; Dalló et al., 2019; Woyann et al., 2020; Jarquin et al., 2020). Since the combination of multivariate techniques is efficient to account for the multicollinearity issue in multi-trait indexes (Rocha et al., 2018; Olivoto et al., 2019; Zuffo et al., 2020), there would seem to be value in an investigation to develop an index that covers the weaknesses of the SH index in which all the traits are selected favorably and with satisfactory gains for the application in breeding programs (Bermudez and Pinheiro, 2020) and biological experiments.

In light of this, we propose a novel multi-trait genotype-ideotype distance index (MGIDI), focused on genotype selection and treatment recommendation based on information of multiple traits. The performance of the proposed index is assessed through Monte Carlo simulations where the success for selecting traits with desired gains is computed for several scenarios varying the number of genotypes and traits evaluated. An application of the index to two real datasets is also presented to illustrate its benefits. The MGIDI index requires a two-way table as input data and provides a row ranking depending on the desired value of columns. Thus, the range of possible applications is spaned to any kind of biological experiment with qualitative or quantitative treatment structure (or even a combination of them), and more than one dependent trait evaluated e.g., Diel et al. (2020); Schwerz et al. (2017); Carvalho et al. (2020); Ferreira et al. (2019). To facilitate the implementation of the MGIDI index in future studies, we provide its implementation in open-source software and guide the user along a gentle coding learning curve.

## 2. Materials and Methods

### 2.1 Theory

The theory of the MGIDI index is centered on four main steps. (i) rescaling the traits so that all have a 0-100 range, (ii) using factor analysis to account four the correlation structure and dimensionality reduction of data, (iii) planning an ideotype based on know/desired values of traits, and (iv) to compute the distance between each genotype to the planned ideotype.

#### 2.1.1 Rescaling the traits

Let **X**_*ij*_ be a two-way table with *i* rows/genotypes/treatments and *j* columns/trait. The rescaled value for the *i* th row and *j* th column (*r***X**_*ij*_) is given by:

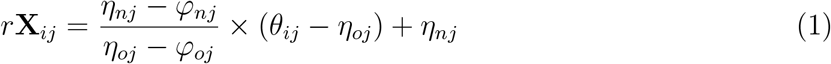

Where *η*_*nj*_ and *φ*_*nj*_ are the new maximum and minimum values for the trait *j* after rescaling, respectively; *η*_*oj*_ and *φ*_*oj*_ are the original maximum and minimum values for the trait *j*, respectively, and *θ*_*ij*_ is the original value for the *j* th trait of the *i* th genotype/treatment. The values for *η*_*nj*_ and *φ*_*nj*_ is chosen as follows. For the traits in which negative gains are desired *η*_*nj*_ = 0 and *φ*_*nj*_ = 100 should be used. For the traits in which positive gains are desired then, *η*_*nj*_ = 100 and *φ*_*nj*_ = 0. In the rescaled two-way table (*r***X**), each column of has a 0-100 range that considers the desired sense of selection (increase or decrease) and maintains the correlation structure of the original set of variables.

#### 2.1.2 Factor analysis

The second step is to compute an exploratory factor analysis to group correlated traits into factors and then estimate the factorial scores for each row/genotype/treatme as follows:

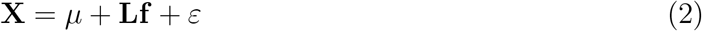

where **X** is a *p ×* 1 vector of observations; *µ* is a *p ×* 1 vector of standardized means; **L** is a *p × f* matrix of factorial loadings; **f** is a *p ×* 1 vector of common factors; and *ε* is a *p ×* 1 vector of residuals, being *p* and *f* the number of traits and common factors retained, respectively. The eigenvalues and eigenvectors are obtained from the correlation matrix of *r***X**_*ij*_. The initial loadings are obtained considering only factors with eigenvalues higher than one. Then, the *varimax* (Kaiser, 1958) rotation criteria is used for the analytic rotation and estimation of final loadings. The scores are then obtained as follows:

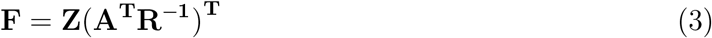

where **F** is a *g × f* matrix with the factorial scores; **Z** is a *g × p* matrix with the (rescaled) standardized means; **A** is a *p × f* matrix of canonical loadings, and **R** is a *p × p* correlation matrix between the traits. *g, f*, and *p* represents the number of rows/genotypes/treatments, factors retained, and analyzed traits, respectively.

#### 2.1.3 Ideotype planning

By definition [Equation (1)], the ideotype has the highest rescaled value (100) for all analyzed traits. Thus, the ideotype can be defined by a 1 *× p* vector **I** such that **I** = [100, 100, *…*, 100]. The scores for **I** is also estimated according to Equation (3).

#### 2.1.4 The MGIDI index

The fourth and last step is the estimation of the multi-trait genotype-ideotype distance index (MGIDI) as follows:

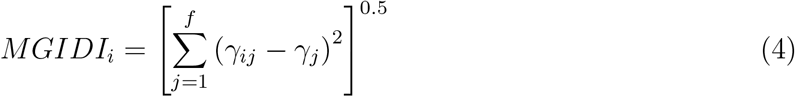

where *MGIDI*_*i*_ is the multi-trait genotype-ideotype distance index for the *i* th row/genotype/treatment; *γ*_*ij*_ is the score of the *i* th row/genotype/treatment in the *j* th factor (*i* = 1, 2, *…, g*; *j* = 1, 2, *…, f*), being *g* and *f* the number of rows/genotypes/treatments and factors, respectively; and *γ*_*j*_ is the *j* th score of the ideotype. The row/genotype/treatment with the lowest MGIDI is then closer to the ideotype and therefore presents desired values for all the *p* traits.

The proportion of the MGIDI index of the *i* th row/genotype/treatment explained by the *j* th factor (*ω*_*ij*_) is used to show the strengths and weaknesses of genotypes/treatments and is computed as:

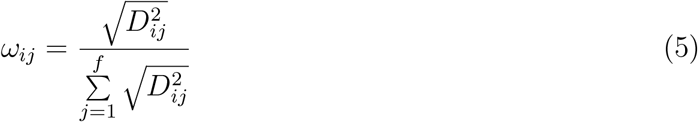

where *D*_*ij*_ is the distance between the *i* th genotype/treatment and the ideotype for the *j* th factor. Low contributions of a factor indicates that the traits within such a factor are close to the ideotype.

### 2.2 Simulation study

We used a Monte Carlo simulation study (N = 500) applied to a simulated dataset (See supplementary Appendix A1.2) to assess the performance of the MGIDI index and compare it with the classical Smith-Hazel (SH) index (Smith, 1936; Hazel, 1943), and the modern FAI-BLUP index (Rocha et al., 2018) in terms of percentage of success in selecting traits with desired gains. Scenarios containing all the combinations of different numbers of genotypes (20 and 200 genotypes) and traits (5, 10, 15, and 20 traits) were planned. For each iteration, genotypes and traits were randomly selected from the simulated data. The ideotype planning was made so that approximately half of the traits had positive desired gains and the other one negative desired gains. A selection intensity of 15% was used. The indexes were computed with the functions `mgidi(), Smith Hazel(), and fai blup()` of the package `metan` (Olivoto and Lúcio, 2020). A benchmark was also performed to assess the computational efficiency of the indexes in different scenarios. The simulations were ran in an Intel(R) Core(TM) i7-9750H CPU @2.6GHz laptop with 16 GB RAM running in multiprocess parallelizes planned with the package `future.apply` (See the code in Supplementary Appendix A1.3).

### 2.3 Real datasets

#### 2.3.1 Dataset 1: Genotype selection in breeding programs

The illustration of the index is made using a replicate-based real dataset coming from a trial with 35 homozygote tropical wheat lines developed by the Wheat Breeding Program of the Federal University of Viçosa (UFV), Viçosa, MG, Brazil in addition to nine commercial cultivar, totaling 44 genotypes. The genotypes were arranged in a randomized complete block design with three replications. Data on 14 agronomic traits (Supplementary Table S1) were assessed. The genotype selection aimed at selecting genotypes with lower values (negative gains) for FLO, PH, EH, and FLH, and higher values (positive gains) for DIS, GY, HW, NSE, NGS, EL, EW, NGE, GME, and HIEP.

Each trait was analyzed using the function `gamem()` of the R package `metan` considering the following mixed-effect model:

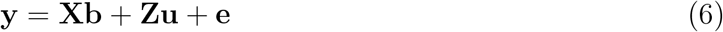

where **y** is an 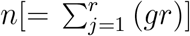 vector of response variable, i.e., the response of the ith genotype in the jth block (*i = 1, 2, …, g; j = 1, 2, …, r* ; **y** = [*y*_11_, *y*_12_, *…, y*_*gr*_]^*′*^); **b** is an 1*×r* vector of unknown and unobservable fixed effects of block **b** = [*γ*_1_, *γ*_2_, *…, γ*_*r*_]^*′*^; **u** is an *m*[= 1 *× g*] vector of unknown and unobservable random effects of genotype **u** = [*α*_1_, *α*_2_, *…, α*_*g*_]^*′*^; **X** is an *n × r* design matrix of 0s and 1s relating **y** to **b**; **Z** is an *n × m* design matrix of 0s and 1s relating **y** to **u**; and **e** is an *n ×* 1 vector of random errors **e** = [*y*_11_, *y*_12_, *…, y*_*gr*_]^*′*^. The variance components generated from the analysis were used to estimate the broad-sense heritability (*h*^2^) on a genotype mean basis as follows:

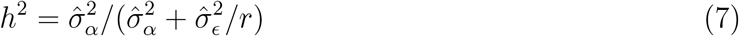

The Best Linear Unbiased Predictors (BLUP) for each genotype and trait were obtained to fill the two-way table described in Equation (1) that was late used to compute the MGIDI, FAI-BLUP, and SH indexes, which were compared in terms of genetic gains. For the SH index, two scenarios were considered. In the first one (SH-1), the SH index was computed with all the 14 traits. In the second one (SH-2), the multicollinearity-generating traits were excluded and the index was computed with the remaining traits. The traits to be excluded were chosen with the function `non collinear vars()` of the package `metan` (Olivoto and Lúcio, 2020) so that there are no VIF great than 10 (Olivoto et al., 2017). For all indexes, it was considered a selection intensity of 15%. To quantify the coincidence of genotype’s selection, we also computed the coincidence index (Hamblin and Zimmermann, 1986) between each pair of indexes.

The predicted genetic gain obtained with the index, *SG*(%), was computed for each trait considering an *α*% selection intensity as follows:

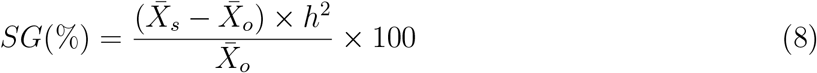

where 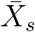 is the mean of the selected genotypes, 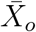 is the mean of the original population and *h*^2^ is the heritability defined in Equation (7).

#### 2.3.2 Dataset 2: Treatment recomendation in agronomic experiments

The second real dataset contains the phenotypic means for the experiment described in Olivoto et al. (2016). The experiment tested the effect of nitrogen splitting at different growth stages and sulfur supplementation on agronomic traits and rheological properties of wheat flour. The MGIDI index is used to rank the treatments according to the desired values of each trait.

Henceforward, the notation S1, S2, … will refer to supplementary figures and/or tables.

## 3. Results

### 3.1 Simulation study

The MGIDI was found to outperform the FAI-BLUP and SH indexes in all simulation scenarios (Fig. 1a). In general, this index presents 74.9% of success in selecting traits with desired gains. FAI-BLUP and SH are successful in 68.5% and 58% of the cases, respectively. The indexes’ performance is dependent on the number of traits and number of genotypes analyzed. For all indexes, the higher success rates occur in datasets with higher number of genotypes and fewer number of traits (Fig. 1a).

**Figure 1.**
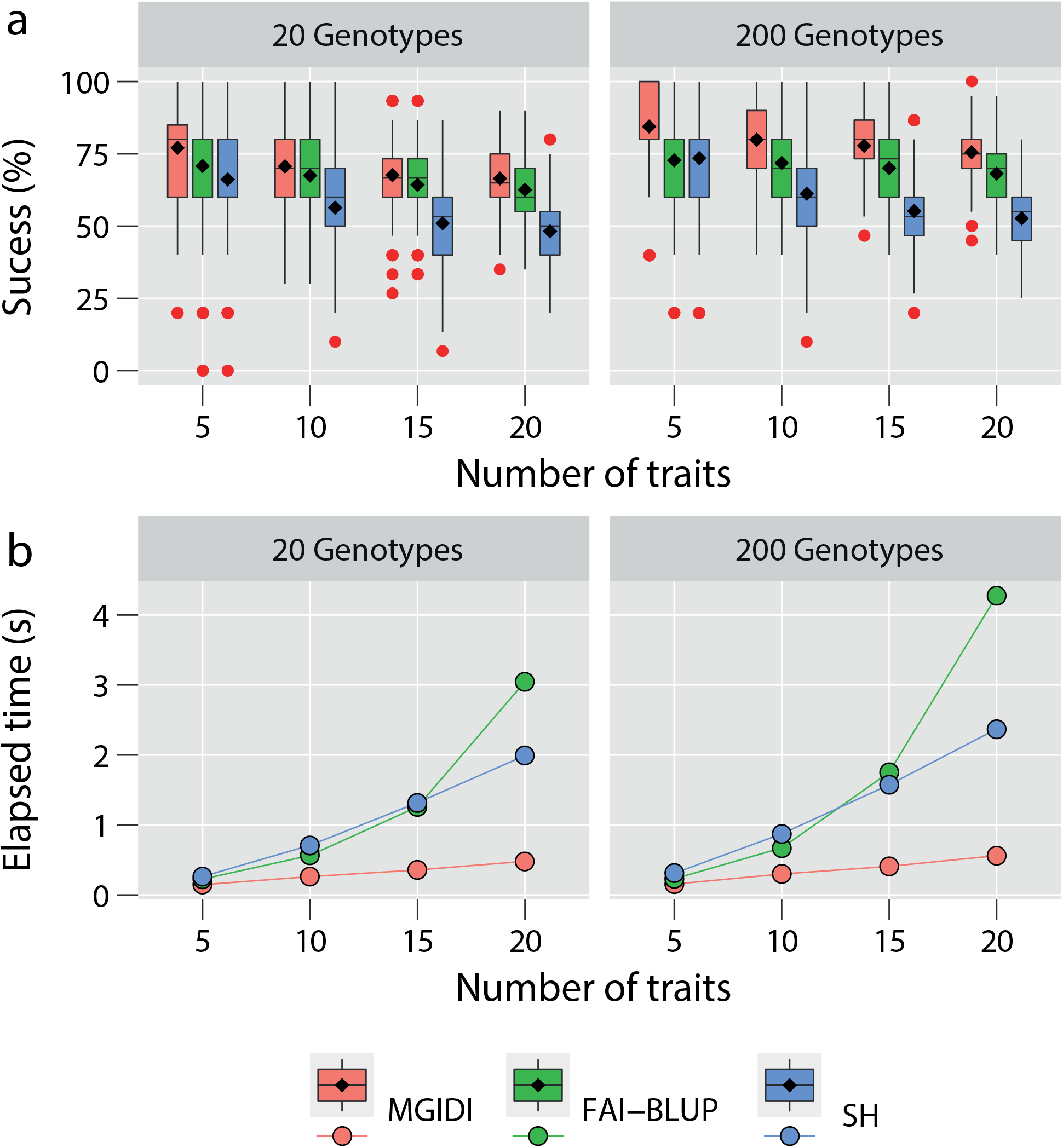
Monte Carlo simulation results (N = 500). The percentage of sucess in selecting traits with desired gains (a) and the average of elapsed time for each step (b) are shown for the indexes MGIDI, FAI-BLUP and Smith-Hazel (SH) in diferent scenarios of number of genotypes (20 and 200 genotypes) and traits (5, 10, 15, and, 20 traits).

Our benchmark results (Fig. 1b) suggests that MGIDI was also more computationally efficient. On average, the computation time to fit the index raised from 0.151 s considering the model with five traits to 0.522 s considering a model with 20 traits. Following this same (clear) linear relationship, it is reasonable to affirm that a model with 50 traits would need nearly one second to fit and that the number of genotypes has no substantial impact on computation time. Both FAI-BLUP and SH indexes needed more than one second to run a model with 15 traits, independently on the number of genotypes. For the SH index, an increasing linear trend of computation time is observed; For the FAI-BLUP, the elapsed time growth exponentially, which may be a worry when several (say, 50 traits) are evaluated.

### 3.2 Dataset 1

Higly significant genotype effects (p *<* 0.005) was observed for all analyzed traits (Supplementary table S2). The broad-sense heritability on a genotype mean basis (*h*^2^) ranged from 0.51 (NGSP) to 0.92 (FLO). High values of heritability (*h*^2^ *>* 0.8) were observed for DIS, FLH, FLO, GMS, HW, SH, and SW (Supplementary table S2), suggesting goods prospects of selection gains for these traits.

Five principal components were retained (Eigenvalue ¿ 1), which explained 87.02% of the total variation among the traits (Supplementary table S3). After the *varimax* rotation, the average communality (h) was 0.87 (HW 0.76 ⩽ *h* ⩽ 0.98 SH), indicating that a high proportion of each variable’s variance was explained by the factors. The 14 traits were grouped into the five factors (FA) as follows: FA1 the spike-related traits (NSS, SL, SW, NGS, and GMS); FA2 the traits related to grain weight (HW and HIS); FA3 the plant-related traits (PH, SH, and FLH); FA4 the traits FLO, DIS, and NGSP; and in the FA5 the GY (Supplementary table S3).

#### 3.2.1 Selected genotypes and coincidence index

The genotypes selected by the MGIDI index were G12, G37, G18, G8, G11, G9, and G32 (Fig. 2). G15 was very close to the cut point (red line that indicates the number of genotypes selected according to the selection pressure), which suggests that this genotype can present interesting features. Thus, the researcher should deserve attention to investigate genotypes that are very close to the cutpoint. Of the seven genotypes selected, the MGIDI index shares only four with the FAI-BLUP index, one with the SH-1 index, and three with the SH-2 index. Only one genotype (G32) was common to all indexes (Supplementary table S4).

**Figure 2.**
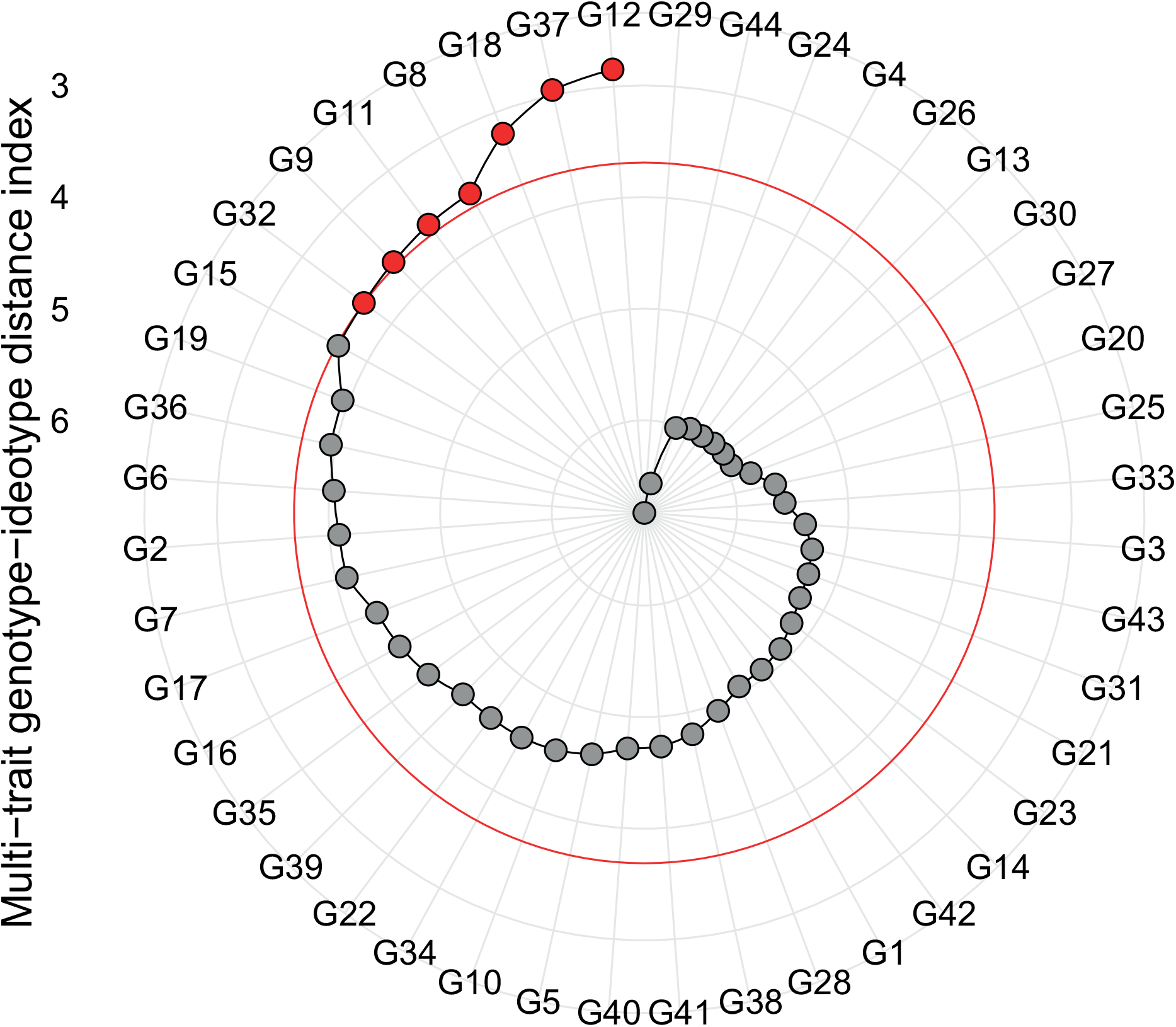
Genotype ranking in ascending order for the MGIDI index. The selected genotypes are shown in red in the electronic version of the article. The circle represents the cutpoint according to the selection pressure. See the ranks for FAI-BLUP, SH-1, and SH-2 indexes as Supplementary Fig. S16.

#### 3.2.2 Predicted selection gains

Table 1 presents the comparison –via selection gains (SG)– between the studied indexes. The number of traits with desired gains was 13, 10, 5, and 9 for MGIDI, FAI-BLUP, SH-1, and SH-2, respectively. This reinforces the results observed in the simulation study, suggestiong that MGIDI was the most efficient index to select genotypes with desired characteristics.

**Table 1.** Predicted genetic gains for the indexes MGIDI, FAI-BLUP, SH-1 and SH-2

| Factor | Trait<sup>a</sup> | Goal | Genetic gain (%) |  |  |  |
| --- | --- | --- | --- | --- | --- | --- |
|  |  |  | MGIDI | FAI-BLUP | SH-1 | SH-2 |
| FA1 | NSS | Increase | 2.22 | -0.56 | -4.05 | -1.88 |
| FA1 | SL | Increase | 3.86 | -0.53 | -4.71 | -2.20 |
| FA1 | SW | Increase | 14.28 | 9.92 | -8.22 | 3.41 |
| FA1 | NGS | Increase | 2.45 | 2.98 | -3.16 | 1.00 |
| FA1 | GMS<sup>b</sup> | Increase | 12.98 | 11.83 | -6.22 | 4.18 |
| FA2 | HW | Increase | 1.44 | 1.66 | 1.11 | 0.35 |
| FA2 | HIS | Increase | -0.55 | 2.13 | 1.71 | 1.08 |
| FA3 | PH | Decrease | -0.07 | 0.09 | 3.86 | 0.10 |
| FA3 | SH<sup>b</sup> | Decrease | -0.74 | 0.19 | 5.49 | 0.49 |
| FA3 | FLH | Decrease | -2.29 | -1.37 | 4.89 | -0.03 |
| FA4 | FLO | Decrease | -2.05 | -2.45 | 4.40 | 1.11 |
| FA4 | DIS | Increase | 8.54 | 4.50 | 20.66 | 11.23 |
| FA4 | NGSP | Increase | 0.46 | 1.94 | 0.41 | 1.43 |
| FA5 | GY | Increase | 3.96 | 4.30 | 4.64 | 7.62 |
| Total (Increase) |  |  | 49.64 | 38.17 | 2.17 | 26.22 |
| Total (Decrease) |  |  | -5.15 | -3.54 | 18.64 | 1.67 |

<sup>a</sup> See Supplementary Table S1 for the full trait names.
<sup>b</sup> Traits removed from SH-2 due to multicollinearity issues.

The only trait with undesired selection gain (*−*0.55%) using the MGIDI index was HIS. Compared to FAI-BLUP, SH-1 and SH-2, MGIDI provided more balanced gains for the analyzed traits. Besides, the MGIDI index provided the higher total gains, i.e., 49.62% for traits that wanted to increase and of *−*5.15% for traits that wanted to decrease (Table 1).

#### 3.2.3 The strengths and weaknesses view

Presented in Fig. 3 is the strengths and weaknesses of genotypes, which are accounted for by the proportion of each factor to the MGIDI index of the genotypes. Let us examine the contribution of FA5 (GY). This factor had the smallest contribution for genotypes G32 and G11. This then indicates that they were the most yielding genotypes among the selected ones (Fig. 4). On the other hand, FA5 had the higher contribution to the MGIDI of G37, suggesting this genotype is poorly productive (Fig. 4). The same interpretation can be extended to other factors.

**Figure 3.**
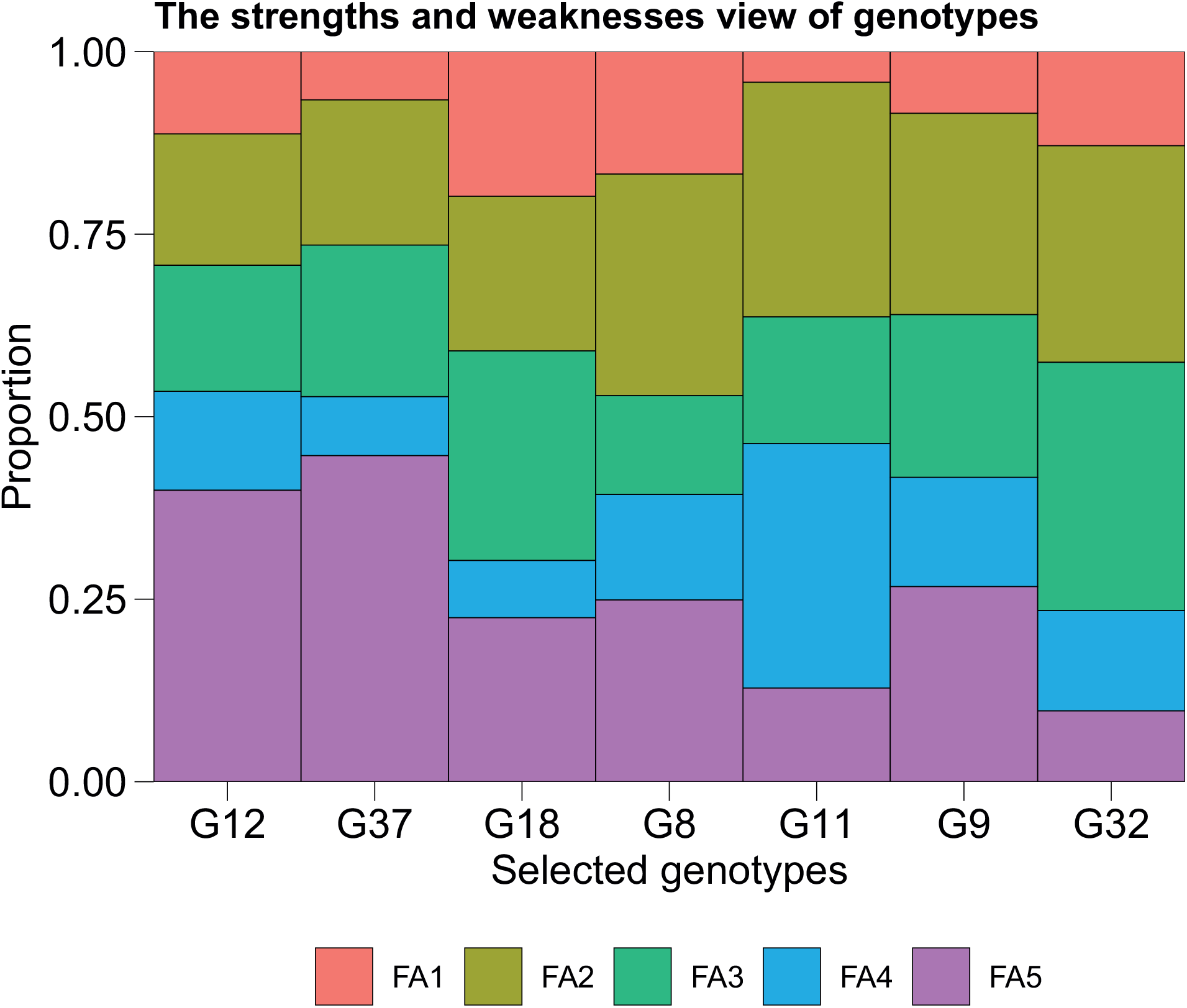
The strengths and weaknesses view of genotypes. The y-axis shows the proportion of each factor on the computed multi-trait genotype-ideotype distance (MGIDI) index of the selected genotypes. The smallest the proportion explained by a factor, the closer the traits within that factor are to the ideotype. The contribution of factors for all the studied genotypes can be found in Supplementary Fig. S2.

**Figure 4.**
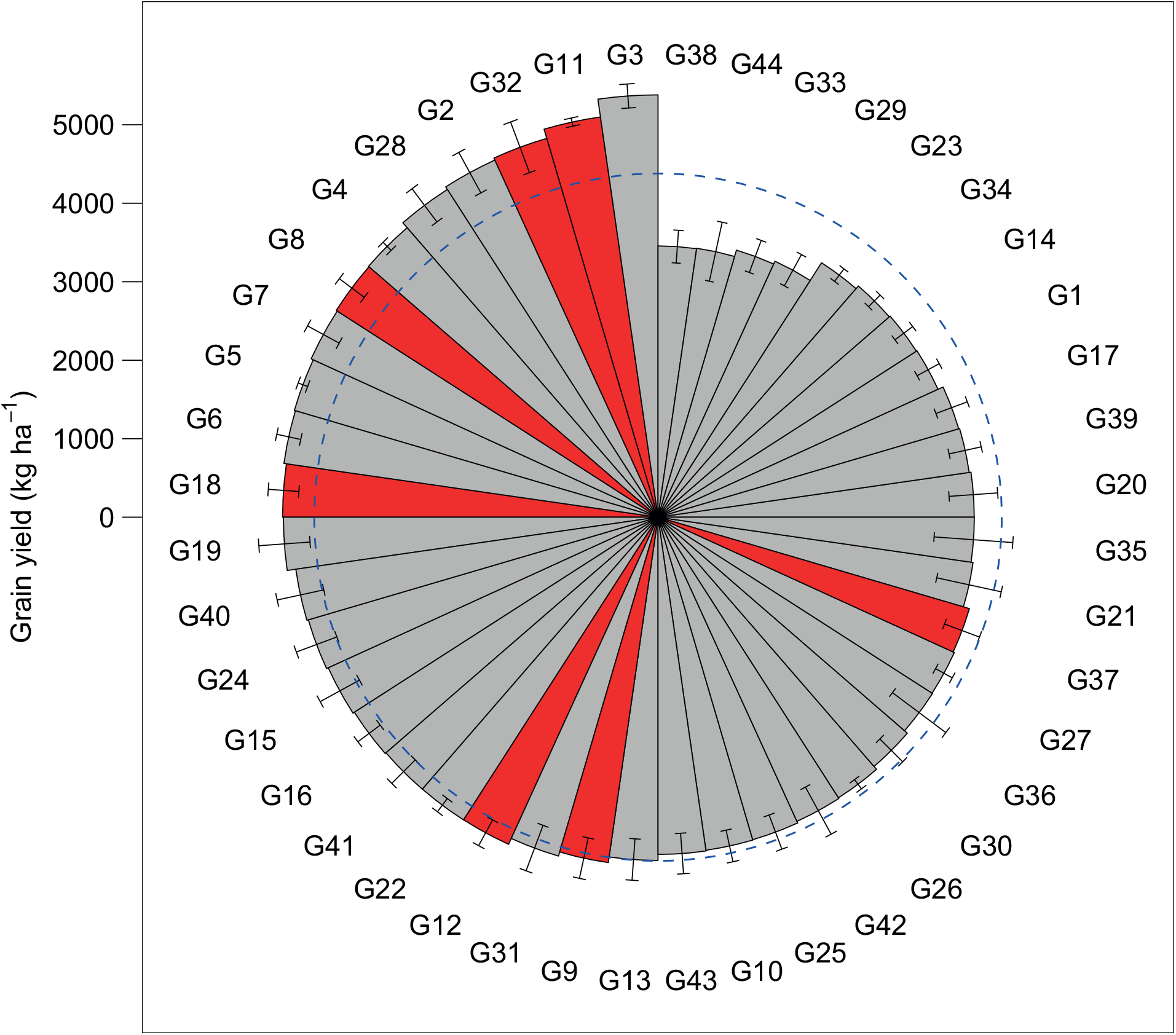
Grain yield for the 44 genotypes tested. The genotypes are sorted in increasing order of grain yield. Genotypes selected by the MGIDI index are shown in red in the electronic version of the article. The dashed line shows the grand mean. Bars show the mean *±*SE. N = 3.

The smallest contributions of FA1 (spike-related traits factor) was observed for G37. Since for all traits in FA1 positive gains are desired, these genotypes should then have (simultaneously) high values for NSS, SL, SW, NGS, and GMS, the traits whithin FA1 (See supplementary Figs S9 and S11-S14). The smallest contributions of FA2 for G12 and G37 (Fig. 3) suggests that these genotypes have high values of HW (Supplementary Fig. S8) and HIS (Supplementary Fig. S15), compared to G11, for example, in which FA2 had high contribution. The smallest contribution of FA3 (plant-related traits) for G8 (Fig. 3) indicates this genotype has a low stature (Supplementary Fig. S4-S6). Since G8 also performed well regarding GY (Fig. 4), this genotype stands out as a potential genitor to obtain segregant populations with few stature and high grain yield. Finally, the smallest contributions of FA4 were observed for G18 and G37, indicating these genotypes have a shorter vegetative period (Supplementary Fig. S3) and above-average values for DIS (Supplementary Fig. S7) and NGSP (Supplementary Fig. S10).

### 3.3 Dataset 2

Supplementary Appendix A1.4.2 shows the application of the MGIDI index to a real dataset with a focus on selecting the best-performing treatments in agronomic experiments. Three factors were retained, explaining 86% of the total variation. The best performing treatment ranked by the MGIDI index (*“WS_30T_40DR_30B”*) combined sulfur supplementation with nitrogen split at tillering, double ring, and booting stages (30%, 40%, and 30% N, respectively). This treatment provided nine of 10 traits with desired gains, which represents a success rate of 90%. The treatment ranking provided by the MGIDI index suggests that nitrogen applied at once in double-ring or in two applications (50% N at the tillering and 50% N at the double ring stage), independently on the sulfur management, are the worse treatments when looking for high grain yield combined with high dough quality.

## 4. Discussion

### 4.1 The theoretical basis of the MGIDI index

The MGIDI index uses the same rescaling procedure as the MTSI index (Olivoto et al., 2019), which is used for genotype selection based on mean performance and stability. This rescaling procedure puts all traits in a 0-100 range, which facilitates the definition of the ideotype. In this case, an ideotype would have a value of 100 for all traits. This is only possible because the rescaling procedure considers the desired direction of selection (increase or decrease the trait values). In future studies, it will be up to the researchers to define the values of *η*_*nj*_ and *φ*_*nj*_ in Equation (1) to rescale the traits.

In a multi-trait framework, the hypothesis is that the traits may be related in some way due to an underlying correlation structure that is unknown beforehand. In the MGIDI index, the factor analysis was used to account for this correlation structure, like in the FAI-BLUP index (Rocha et al., 2018). This was only possible because the rescaling procedure kept the original correlation structure of the data, allowing to plan an ideotype easily. Thus, it was possible to determine in how many latent variables the original set of traits could be reduced and the relationships among traits within latent variables. In dataset 1, the 14 original traits were reduced into five final latent variables (factors) that explained ∼87% of the original data. This dimensional reduction makes it simpler the interpretation and decision making by practitioners. Because factor analysis produces orthogonal axes among final factors, it was possible to obtain genotype or treatment’s scores free from multicollinearity. Finally, the distance from a given genotype/treatment to the planned ideotype could be computed using the Euclidean distance. So, in contrast to conclusions of recent studies (Bermudez and Pinheiro, 2020), we have shown here that factor analysis is an efficient technique to establish an index in which most of the traits are selected favorably.

In our simulation study, the MGIDI index was found to outperform the FAI-BLUP and SH indexes in selecting traits with desired gains. In addition to providing more success in selecting traits with desired gains, the MGIDI index was computationally more efficient. In the FAI-BLUP, for example, the increase in time computing growths exponentially, since the number of ideotypes to be planned is the *n*th potency of retained factors (Rocha et al., 2018). For example, the number of analyses to be computed would range from 32,768 with 15 factors to 1,048,576 with 20 factors retained. We need to point out that no information regarding the number of retained factors was generated in the simulation study, but is expected that for a dataset with a low correlation among the traits the number of retained factors be higher, increasing considerably the needed time to compute the index.

Correlated data are common in breeding experiments (Nardino et al., 2016). Consequently, the multicollinearity issue (Graham, 2003; Olivoto et al., 2017), is an obstacle faced by the SH index. Since the SH index requires inverting a phenotypic covariance matrix among traits, the presence of highly correlated traits can result in either biased index coefficients –since the phenotypic (co)variance matrix is not optimally conditioned– or in an infinite number of solutions if such a matrix is not positive definite. Even though non-collinear traits can be selected easily using the function `non collinear vars()` of the R package `metan` (Olivoto and Lúcio, 2020) –which would facilitate the implementation of the SH index in future studies– we strongly suggest the use of the MGIDI index, since it takes multicollinearity into account, provides a high rate of success in selecting traits with desired gains, and is more computationally efficient then FAI-BLUP index.

### 4.2 The MGIDI in practice

The MGIDI index was found to have many practical applications since it allows a unique and easy-to-interpret selection process. Besides dealing with collinear traits, the MGIDI index doesn’t require the use of economic weights such as in the SH index, in which one can predict genetic and economic gains for various possible combinations of genetic parameters and assumed economic weights (Bizari et al., 2017; Burdon and Li, 2019). In our study, the MGIDI index was found to outperform the FAI-BLUP and SH indexes since it provided more balanced gains. Thus, MGIDI can help breeders to guarantee long-term gains in primary traits (e.g., grain yield) without jeopardizing genetic gains of secondary traits (e.g., plant height).

Functions to compute the MGIDI, FAI-BLUP, and SH indexes in R software have been elaborated and implemented in the R package `metan` (Olivoto and Lúcio, 2020). These functions will make it easier the implementation of these indexes in future studies. The researcher uses a model fitted with the functions `gafem()` (fixed model) or `gamem()` (mixed model) as the input data in the function `mgidi()`, plans the ideotype by declaring which traits are desired to increase or decrease and the function will take care of the details.

The *“strengths and weaknesses view”*, i.e., the proportion of the MGIDI index explained by each factor (Fig. 3) is an important tool to identify the strengths and weaknesses of genotypes or treatments. From a breeder point of view, it is possible to identify in the selected genotypes –or even in the non-selected ones (Supplementary Fig. S2)– traits or groups of traits that need to be improved. For example, the low contribution of FA5 in G11 (Fig. 3) indicates that this genotype is highly productive (Fig. 4) but was the selected genotype with lesser hectoliter weight (Supplementary Fig. S8) and lesser harvest index of the spike (Supplementary Fig. S15), which can be inferred due to the high contribution of FA2 (Fig. 3). G11 was also the most health (Supplementary Fig. S7) and with the longer vegetative period (Supplementary Fig. S3) among the selected genotypes, which is explained by the high contribution of FA4 for this genotype (Fig. 3). We may use then these contributions to chose possible genitors for future crossings. In our example, G8, G11, and G18 could be included in a crossing block aiming at obtaining plants with high production and healthy (G11), with a shorter vegetative period (G18), and with a lesser stature (G8). From an agronomist point of view, the MGIDI index can help practitioners to make better strategic decisions for treatment recommendations. Compared to state-of-the-art methods, this approach is graphical, objective, effective, and straightforward to identify the strengths and weaknesses of genotypes or treatments based on multiple traits.

### 4.3 A step-by-step guide for future studies

Our Web Appendix A provides the codes used in this article that can be easily adapted in future studies. There have been two main ways to compute the MGIDI index. The first is by using a two-way table of the genotype/treatment-vs-trait mean (Supplementary Appendix A1.5.1) In this case, since there is no information regarding the heritability of the traits, only the selection differentials are computed. The second one is by using raw data (Supplementary Appendix A1.5.2). In this case, the MGIDI index can be computed from models previously fitted with the functions `gafem()` (fixed-effect models) and `gamem()` (mixed-effect model), which was we have done in this article. When a fitted model is used as input data, the heritability is extracted internally, allowing the computation of selection gains automatically.

Another useful function of the `metan` (Olivoto and Lúcio, 2020) package that can facilitate the implementation of the multi-trait indexes in future studies is `coincidence index()` (Supplementary Appendix A1.5.3). This function will make it easy —and we strongly suggest – the comparison of the indexes in terms of predicted genetic gains in future studies.

## Acknowledgements

We thank the graduate post-graduate students of the Federal University of Viçosa, who helped in the data collection and typing. We also

## References

Bermudez, F. and Pinheiro, J. B. (2020). Selection to high productivity and stink bugs resistance by multivariate data analyses in soybean. Bragantia 79, 250–259.

Bhering, L. L., Laviola, B. G., Salgado, C. C., Sanchez, C. F. B., Rosado, T. B., and Alves, A. A. (2012). Genetic gains in physic nut using selection indexes. Pesquisa Agropecuaria Brasileira 47, 402–408.

Bizari, E. H., Val, B. H. P., Pereira, E. d. M., Mauro, A. O. D., and Unêda-Trevisoli, S. H. (2017). Selection indices for agronomic traits in segregating populations of soybean. Revista Ciencia Agronomica 48, 110–117.

Burdon, R. D. and Li, Y. (2019). Genotype-environment interaction involving site differences in expression of genetic variation along with genotypic rank changes: simulations of economic significance. Tree Genetics & Genomes 15, 1–10.

Carvalho, I. R., Szareski, V. J., da Silva, J. A. G., Nunes, A. C. P., da Rosa, T. C., Barbosa, M. H., Magano, D. A., Conte, G. G., Caron, B. O., and de Souza, V. Q. (2020). Multivariate best linear unbiased predictor as a tool to improve multi-trait selection in sugarcane. Pesquisa Agropecuária Brasileira 55,.

Cerón-Rojas, J. J. and Crossa, J. (2018). Linear selection indices in modern plant breeding. Springer International Publishing.

Dalló, S. C., Zdziarski, A. D., Woyann, L. G., Milioli, A. S., Zanella, R., Conte, J., and Benin, G. (2019). Across year and year-by-year GGE biplot analysis to evaluate soybean performance and stability in multi-environment trials. Euphytica 215, 1–12.

Diel, M. I., Lućio, A. D., Olivoto, T., Pinheiro, M. V. M., Krysczun, D. K., Sari, B. G., and Schmidt, D. (2020). Repeatability coefficients and number of measurements for evaluating traits in strawberry. Acta Scientiarum. Agronomy 42, e43357.

Donald, C. M. (1968). The breeding of crop ideotypes. Euphytica 17, 385–403.

Ferreira, C. D., Ziegler, V., Schwanz Goebel, J. T., Hoffmann, J. F., Carvalho, I. R., Chaves, F. C., and De Oliveira, M. (2019). Changes in Phenolic Acid and Isoflavone Contents during Soybean Drying and Storage. Journal of Agricultural and Food Chemistry 67, 1146–1155.

Graham, M. H. (2003). Confronting Multicollinearity in Ecological Multiple Regression. Ecology 84, 2809–2815.

Hamblin, J. and Zimmermann, M. J. d. O. (1986). Breeding Common Bean for Yield in Mixtures. In Plant Breeding Reviews, pages 245–272. John Wiley & Sons, Inc., Hoboken, NJ, USA.

Hazel, L. N. (1943). The Genetic Basis for Constructing Selection Indexes. Genetics 28, 476–90.

Jahufer, M. Z. Z. and Casler, M. D. (2015). Application of the Smith-Hazel Selection Index for Improving Biomass Yield and Quality of Switchgrass. Crop Science 55, 1212–1222.

Jarquin, D., Howard, R., Crossa, J., Beyene, Y., Gowda, M., Martini, J. W. R., Covarrubias-Pazaran, G., Burgueño, J., Pacheco, A., Grondona, M., Wimmer, V., and Prasanna, B. M. (2020). Genomic Prediction Enhanced Sparse Testing for Multi-environment Trials. G3: Genes, Genomes, Genetics page g3.401349.2020.

Kaiser, H. F. (1958). The varimax criterion for analytic rotation in factor analysis. Psychometrika 23, 187–200.

Nardino, M., de Souza, V. Q., Baretta, D., Konflanz, V. A., Carvalho, I. R., Follmann, D. N., and Caron, B. O. (2016). Association of secondary traits with yield in maize F1’s. Ciência Rural 46, 776–782.

Olivoto, T., Carvalho, I. R., Nardino, M., Ferrari, M., de Pelegrin, A. J., Follmann, D. N., Gutkoski, L. C., and de Souza, V. Q. (2016). Sulfur and nitrogen effects on industrial quality and grain yield of wheat. Revista de Ciências Agroveterinárias 15, 24–33.

Olivoto, T. and Lućio, A. D. (2020). metan: An R package for multi-environment trial analysis. Methods in Ecology and Evolution 11, 783–789.

Olivoto, T., Lućio, A. D., Silva, J. A., Marchioro, V. S., Souza, V. Q., and Jost, E. (2019). Mean Performance and Stability in Multi-Environment Trials I: Combining Features of AMMI and BLUP Techniques. Agronomy Journal 111, 2949–2960.

Olivoto, T., Lućio, A. D. C., Silva, J. A. G., Sari, B. G., and Diel, M. I. (2019). Mean Performance and Stability in Multi-Environment Trials II: Selection Based on Multiple Traits. Agronomy Journal 111, 2961–2969.

Olivoto, T., Souza, V. Q., Nardino, M., Carvalho, I. R., Ferrari, M., Pelegrin, A. J., Szareski, V. J., and Schmidt, D. (2017). Multicollinearity in path analysis: a simple method to reduce its effects. Agronomy Journal 109, 131–142.

Rocha, J. R. d. A. S. d. C., Machado, J. C., and Carneiro, P. C. S. (2018). Multitrait index based on factor analysis and ideotype-design: proposal and application on elephant grass breeding for bioenergy. GCB Bioenergy 10, 52–60.

Schwerz, F., Sgarbossa, J., Olivoto, T., Elli, E. F., Aguiar, A., Caron, B., and Schmidt, D. (2017). Solar radiation levels modify the growth traits and bromatological composition of cichorium intybus. Advances in Horticultural Science 31, 257–265.

Smith, H. (1936). A discriminant function for plant selection. Annals of Eugenics 7, 240–250.

Woyann, L. G., Meira, D., Matei, G., Zdziarski, A. D., Dallacorte, L. V., Madella, L. A., and Benin, G. (2020). Selection indexes based on linear-bilinear models applied to soybean breeding. Agronomy Journal 112, 175–182.

Zuffo, A. M., Steiner, F., Aguilera, J. G., Teodoro, P. E., Teodoro, L. P. R., and Busch, A. (2020). Multi-trait stability index: A tool for simultaneous selection of soya bean genotypes in drought and saline stress. Journal of Agronomy and Crop Science 00, 1–8.

