## Supplementary for "MGIDI: towards an effective multivariate selection in biological experiments"

### Supporting Informatin for 'MGIDI: a novel multi-trait index for genotype selection in plant breeding'

**3 Appendix C** **41**

**References** **45**

### Getting started

This supplementary material has three main sections. In the first one (Appendix A), we present the R datasets, the packages, and all R codes needed to reproduce the examples described in the paper. The second section (Appendix B), contains the supplementary figures, and the last one (Appendix C) contains the supplementary tables.

### 1 Appendix A

#### 1.1 R pakages

To reproduce the examples of this material, the R packages `metan` (Olivoto & Lúcio, 2020) `tidyverse` (Wickham et al., 2019), and `future.apply` (Bengtsson, 2020) are needed. If you don't have these packages yet, install them by running `install.packages("metan")`, `install.packages("tidyverse")`, and `install.packages("future.apply")`, respectively, in the R console. Then, load the packages with:

```
library(metan)
library(tidyverse)
library(future.apply)
```

#### 1.2 Datasets

In this section we provide the access to the three datasets used in the study.

1. The first dataset is used to illustrate the index and contains data on 14 traits eavaluated in 44 wheat genotypes. For more details see the The supplementary Table S1 in Apendix C.

```
data <-
  read.csv("https://bit.ly/2ZOA7FL", sep = ";") %>%
  to_factor(GEN, BLOCK) # two first columns as factor
str(data)
# 'data.frame': 132 obs. of  16 variables:
#  $ GEN  : Factor w/ 44 levels "G1","G10","G11",..: 1 1 1 12 12 12 23 23 23 34 ...
#  $ BLOCK: Factor w/ 3 levels "1","2","3": 1 2 3 1 2 3 1 2 3 1 ...
#  $ FLO  : int  67 67 67 66 67 67 69 69 69 67 ...
#  $ PH   : num  88.3 93.5 88.6 87 84.7 79.1 80.5 86.6 88.1 90.2 ...
#  $ SH   : num  80.3 84.1 81.6 80.7 78.1 70.7 74.2 78.8 80.8 82.9 ...
#  $ FLH  : num  65.6 68.7 65.5 67.5 63.1 59.9 59.8 61.7 65.7 64.1 ...
#  $ DIS  : int  4 4 3 5 5 4 5 5 5 2 ...
#  $ GY   : num  4223 3867 3676 4800 4626 ...
#  $ HW   : num  77.4 77.1 76 80.8 77.6 ...
#  $ NSS  : num  12.6 15.8 15.3 15.2 13.8 15.1 13.2 14.3 13.7 15.5 ...
```

```
#  $ NGSP : num  3.13 2.52 2.73 2.49 2.51 2.7 2.6 2.38 2.27 2 ...
#  $ SL   : num  8 9.4 7 6.3 6.6 8.4 6.3 7.8 7.3 7.3 ...
#  $ SW   : num  1.96 2.02 1.68 1.86 1.57 1.9 1.64 1.78 1.39 1.42 ...
#  $ NGS  : num  39.3 39.7 41.7 37.9 34.4 40.7 34.3 33.9 31.8 31.2 ...
#  $ GMS  : num  1.45 1.5 1.27 1.49 1.28 1.45 1.3 1.33 1.16 1.07 ...
#  $ HIS  : num  73.8 74.3 75.6 80.2 81.4 ...
```

2. The second dataset contains the phenotypic means for the experiment described in (Olivoto et al., 2016). The experiment tested the effect of nitrogen splitting at different stages (DR, double ring; T, tillering; and B, booting stage) and sulfur application (with and without sulfur) on agronomic and rheological characteristics. The data contains the following columns: TRAT: the treatment column. P: Tenacity (mm) L: Extensibility (mm) PL: P/L ratio TIL: number of tillers SSM: spikes per square meter GY: Grain yield (kg/ha) HW: Hectoliter weight W: Gluten strength (W $\times 10^{-4}$ J) GLU: Gluten content (%) PROT: Protein content (%).

```
data_ns <-
  read.csv("https://bit.ly/3jKx8Jo", sep = ";")
str(data_ns)
# 'data.frame': 8 obs. of  11 variables:
#  $ TRAT: chr  "WS_100DR" "WS_30T_40DR_30B" "WS_50T_50DR" "WS_50DR_50B" ...
#  $ P   : num  91.5 96.8 93.5 96.8 103.5 ...
#  $ PL  : num  0.99 1.14 1.32 1.12 1.31 1.1 1.38 1.29
#  $ L   : num  93.2 85.2 71.5 86.8 79 ...
#  $ TIL : num  3.56 5.48 6.55 4.47 3.53 5.42 5.33 4.42
#  $ SSM : num  582 794 702 626 550 ...
#  $ GY  : num  5301 6459 6079 5697 5485 ...
#  $ HW  : num  79.2 79.2 79.2 79.5 78.8 ...
#  $ W   : num  261 264 224 263 270 ...
#  $ GLU : num  30.1 29.9 27.9 29.6 27.6 ...
#  $ PROT: num  11.8 11.9 12.6 12.6 12.7 ...
```

3. The third dataset is a simulated dataset used to run the Monte Carlo simulations.

```
data_simula <-
  read.csv("https://bit.ly/2GoiJnS", sep = ",") %>%
  to_factor(GEN, REP) # two first columns as factor

str(data_simula)
# 'data.frame': 3000 obs. of  34 variables:
#  $ GEN: Factor w/ 1000 levels "1","2","3","4",..: 1 1 1 2 2 2 3 3 3 4 ...
#  $ REP: Factor w/ 3 levels "1","2","3": 1 2 3 1 2 3 1 2 3 1 ...
#  $ V1 : num  92.9 88.1 93.4 89.9 91.6 ...
```

```
#  $ V2 : num  0.71 0.67 0.61 0.59 0.64 0.6 0.67 0.63 0.7 0.63 ...
#  $ V3 : num  2633 2633 -735 -2271 1455 ...
#  $ V4 : num  412 412 266 368 355 ...
#  $ V5 : num  12811 12811 12828 12740 12802 ...
#  $ V6 : num  208 208 210 273 253 ...
#  $ V7 : num  471 471 460 511 452 ...
#  $ V8 : num  454 454 472 380 444 ...
#  $ V9 : num  273 273 260 254 237 ...
#  $ V10: num  118 118 144 165 133 ...
#  $ V11: num  519 519 494 434 474 ...
#  $ V12: num  674 674 665 584 683 ...
#  $ V13: num  343.5 137.6 11.4 206.7 258.6 ...
#  $ V14: num  497 497 490 350 472 ...
#  $ V15: num  48.7 48.7 60.6 54.1 41.1 ...
#  $ V16: num  181 181 228 143 122 ...
#  $ V17: num  337 337 327 293 323 ...
#  $ V18: num  139 139 136 128 149 ...
#  $ V19: num  104 104 108 137 107 ...
#  $ V20: num  249 249 259 314 266 ...
#  $ V21: num  433 433 387 292 318 ...
#  $ V22: num  178 185 142 198 218 ...
#  $ V23: num  160 166 161 158 153 ...
#  $ V24: num  495 495 471 496 499 ...
#  $ V25: num  3376 3376 3889 5779 4901 ...
#  $ V26: num  -693 -668 -588 -563 -626 ...
#  $ V27: num  -282 -351 -198 -157 -263 ...
#  $ V28: num  85.8 -17.3 124.9 159.9 66.8 ...
#  $ V29: num  424 314 409 429 373 ...
#  $ V30: num  717 663 623 608 645 ...
#  $ V31: num  101881 35316 41776 66198 -250273 ...
#  $ V32: num  507652 494406 488704 355738 412663 ...
```

#### 1.3 Simulation study

Here, we provide the function `run_simula()` to run the Monte Carlo simulation with different number of genotypes and traits.

```
run_simula <- function(df, # The data set (data_simula)
                       nvar, # The number of traits in the simulation
                       ngen,  # The number of genotypes in the simulation
                       index, # The index (one of 'fai', 'mgidi' or 'smith')
                       nboot = 500){ # The number of replications
  if(!index %in% c("smigh", "mgidi", "fai")){
    stop("Invalid index. Use 'fai', 'mgidi' or 'smith'")
```

```
}
# Helper function
sel_gain <- function(df,
                     ngen,
                     index,
                     nvar,
                     ...){
  vars <- names(df[3:ncol(df)])
  ideotype <- c(rep("h", round(nvar / 2)), rep("l", nvar - round(nvar / 2)))
  ideotype_n <- sample(ideotype, nvar)
  # create a sample data set
    datasel <-
    subset(df, GEN %in% sample(unique(df$GEN), ngen, replace = FALSE)) %>%
    select_cols(GEN, REP, any_of(sample(vars, nvar))) %>%
    droplevels()
    # Fit the mixed model
  mod <- gamem(datasel, GEN, REP, everything(), verbose = FALSE)
  if(index == "mgidi"){
    mgid_index <- mgidi(mod,
                        ideotype = if_else(ideotype_n == "h", "h", "l"),
                        use_data = "pheno",# Use phenotypic mean instead BLUPs
                        # to avoid error in FAI-BLUP with BLUPs = 0
                        verbose = FALSE)
    return(mgid_index$sel_dif %>%
             means_by() %>% # Compute the mean of selection success
             select(goal) %>% # select the column goal
             rename(SUCESS = goal))
  }
  if(index == "fai"){
    fai_index <- fai_blup(mod,
                          use_data = "pheno",#Use phenotypic data instead BLUPs
                          # to avoid error in FAI-BLUP with BLUPs = 0
                          DI = if_else(ideotype_n == "h", "max", "min"),
                          UI = if_else(ideotype_n == "h", "min", "max"),
                          verbose = FALSE)
    return(fai_index$selection_diferential$ID1 %>%
             means_by() %>% # Compute the mean of selection success
             select(goal) %>% # select the column goal
             rename(SUCESS = goal))
  }
  if(index == "smith"){
    smith_index <- Smith_Hazel(mod, weights = if_else(ideotype_n == "h", 1, -1))
    return(smith_index$sel_dif %>%
             means_by() %>% # Compute the mean of selection success
```

```
               select(goal) %>% # select the column goal
               rename(SUCESS = goal))
    }
  }
  # Make combinations of gen and trait
  comb <- data.frame(
    expand.grid(GEN = ngen,
                TRAITS = nvar)
  )
  temp <- list()
  elapsed <- list()
  for(i in 1:nrow(comb)){
    code <- paste(comb[i, 1], comb[i, 2])
    print(code)
    elapsed[[code]] <- system.time( # benchmark
      temp[[code]] <-
        do.call(rbind,
                future_lapply(1:nboot, # apply sel_gain nboot times
                              FUN = sel_gain,
                              df = df,
                              index = index,
                              nvar = comb[i, 2],
                              ngen = comb[i, 1])) %>%
        add_cols(INDEX = index, GEN = comb[i, 1],  .before  = 1)
    )
  }
  # bind the rows and get the results for sucess and elapsed time
return(
  list(
    gains = do.call(rbind, lapply(temp, function(x){x}))%>%
      as.data.frame() %>%
      rownames_to_column() %>%
      separate(rowname, into = c("GEN", "TRAIT"), extra = "drop"),
    elapsed = do.call(rbind, lapply(elapsed, function(x){x})) %>%
      as.data.frame() %>%
      rownames_to_column() %>%
      separate(rowname, into = c("GEN", "TRAIT"), extra = "drop")
  )
)
}

# Run the simulations
library(future.apply) # Run in multiprocess (for windows users)
plan(multiprocess)
```

```
# mgidi index
mgidi_ind <-
run_simula(data_simula,
           nvar = c(5, 10, 15, 20),
           ngen = c(20, 200),
           index = "mgidi",
           nboot = 500)
# smith index
smith_ind <-
  run_simula(data_simula,
             nvar = c(5, 10, 15, 20),
             ngen = c(20, 200),
             index = "smith",
             nboot = 500)
# fai index
fai_ind <-
  run_simula(data_simula,
             nvar = c(5, 10, 15, 20),
             ngen = c(20, 200),
             index = "fai",
             nboot = 500)
```

#### 1.4 Real dataset

##### 1.4.1 Dataset 1 - Wheat genotypes

The function `gamem()` is used to fit the mixed-effect model considering genotype as random effect. To analyze all the numeric variables of the data set at once we use `everything()` in the argument `resp`. This will allow to use the object `mod` as input argument in functions to compute the MGIDI, FAI-BLUP and Smith-Hazel indexes.

```
# Mixed-effect model for all variables in the data set
mod <- gamem(data,
             gen = GEN,
             rep = BLOCK,
             resp = everything())
# Method: REML/BLUP
# Random effects: GEN
# Fixed effects: REP
# Denominador DF: Satterthwaite's method
# ---------------------------------------------------------------------------
# P-values for Likelihood Ratio Test of the analyzed traits
# ---------------------------------------------------------------------------
#     model       FLO        PH        SH       FLH       DIS        GY        HW
```

```
#  Complete        NA        NA        NA        NA        NA        NA        NA
#  Genotype 5.31e-23 1.66e-09 1.47e-11 6.21e-12 1.36e-18 5.71e-07 2.19e-10
#       NSS     NGSP        SL       SW       NGS       GMS       HIS
#        NA       NA        NA       NA        NA        NA        NA
#  1.15e-08 0.00465 1.45e-05 1.3e-12 2.89e-05 5.24e-11 1.08e-09
# ---------------------------------------------------------------------------------
# All variables with significant (p < 0.05) genotype effect
```

```
lrt <- gmd(mod, "lrt")
# Class of the model: gamem
# Variable extracted: lrt
lrt
# # A tibble: 14 x 8
#    VAR   model     npar  logLik   AIC   LRT    Df `Pr(>Chisq)`
#    <chr> <chr>    <dbl>   <dbl> <dbl> <dbl> <dbl>        <dbl>
#  1 FLO   Genotype     4  -367.   742.  97.5      1     5.31e-23
#  2 PH    Genotype     4  -410.   828.  36.3      1     1.66e- 9
#  3 SH    Genotype     4  -416.   840.  45.6      1     1.47e-11
#  4 FLH   Genotype     4  -402.   811.  47.3      1     6.21e-12
#  5 DIS   Genotype     4  -207.   421.  77.5      1     1.36e-18
#  6 GY    Genotype     4 -1011.  2029.  25.0      1     5.71e- 7
#  7 HW    Genotype     4  -321.   650.  40.3      1     2.19e-10
#  8 NSS   Genotype     4  -238.   484.  32.6      1     1.15e- 8
#  9 NGSP  Genotype     4   -38.7   85.4  8.01     1     4.65e- 3
# 10 SL    Genotype     4  -196.   401.  18.8      1     1.45e- 5
# 11 SW    Genotype     4   -74.2  156.  50.3      1     1.30e-12
# 12 NGS   Genotype     4  -405.   818.  17.5      1     2.89e- 5
# 13 GMS   Genotype     4   -38.0   84.0 43.1      1     5.24e-11
# 14 HIS   Genotype     4  -365.   738.  37.2      1     1.08e- 9
```

###### 1.4.1.1 LRT

```
vcomp <- gmd(mod, what = "vcomp")
# Class of the model: gamem
# Variable extracted: vcomp
vcomp
# # A tibble: 2 x 15
#   Group   FLO    PH    SH   FLH   DIS     GY    HW   NSS    NGSP    SL     SW
#   <chr> <dbl> <dbl> <dbl> <dbl> <dbl>  <dbl> <dbl> <dbl>  <dbl> <dbl>  <dbl>
```

```
# 1 GEN    12.7   16.9  20.4  16.6 0.980 1.58e5  4.47  1.12 0.0256 0.452 0.107
# 2 Resi~   3.16  14.0  13.4  10.5 0.340 1.85e5  3.33  1.03 0.0721 0.674 0.0628
# # ... with 3 more variables: NGS <dbl>, GMS <dbl>, HIS <dbl>

# Create a plot
plot(mod, type = "vcomp") +
  geom_hline(yintercept = 0.5, linetype = 2)
```

###### 1.4.1.2 Variance components

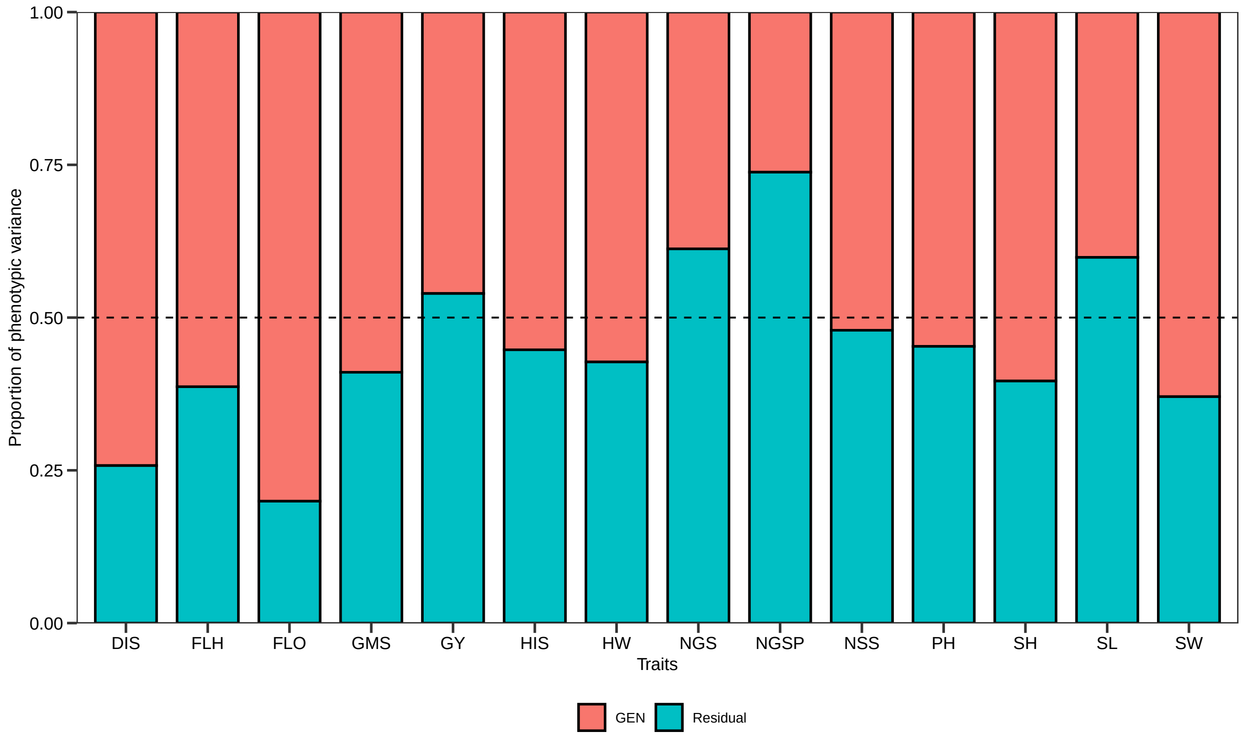

```
gen_par <- gmd(mod) # what = "genpar" is the default value
# Class of the model: gamem
# Variable extracted: genpar
gen_par
# # A tibble: 11 x 15
#    Parameters    FLO     PH     SH    FLH    DIS      GY     HW    NSS    NGSP
#    <chr>       <dbl>  <dbl>  <dbl>  <dbl>  <dbl>   <dbl>  <dbl>  <dbl>   <dbl>
#  1 Gen_var     12.7   16.9   20.4   16.6    0.980 1.58e+5  4.47   1.12   0.0256
#  2 Gen (%)     80.1   54.7   60.4   61.3   74.2   4.61e+1 57.3   52.1   26.2
#  3 Res_var      3.16  14.0   13.4   10.5    0.340 1.85e+5  3.33   1.03   0.0721
```

```
#  4 Res (%)     19.9   45.3   39.6   38.7   25.8   5.39e+1 42.7   47.9   73.8
#  5 Phen_var    15.9   30.9   33.9   27.2    1.32  3.42e+5  7.80   2.15   0.0977
#  6 H2           0.801  0.547  0.604  0.613  0.742 4.61e-1  0.573  0.521  0.262
#  7 h2mg         0.923  0.784  0.821  0.826  0.896 7.19e-1  0.801  0.765  0.516
#  8 Accuracy     0.961  0.885  0.906  0.909  0.947 8.48e-1  0.895  0.875  0.718
#  9 CVg          5.89   4.75   5.81   6.75  34.8   9.07e+0  2.77   6.80   6.11
# 10 CVr          2.94   4.32   4.71   5.36  20.5   9.81e+0  2.39   6.53  10.3
# 11 CV ratio     2.00   1.10   1.23   1.26   1.70  9.24e-1  1.16   1.04   0.596
# # ... with 5 more variables: SL <dbl>, SW <dbl>, NGS <dbl>, GMS <dbl>,
# #   HIS <dbl>
```

###### 1.4.1.3 Genetic parameters

```
gcov <- gmd(mod, what = "gcov")
# Class of the model: gamem
# Variable extracted: gcov
round_cols(gcov, digits = 0) # to fit in the page width
#      DIS FLH FLO GMS     GY HIS HW NGS NGSP NSS  PH  SH  SL SW
# DIS    1   0   2   0     89   1  1  -2    0   0   0   0   0  0
# FLH    0  17   9  -1    120   3  1  -7    0  -1  14  16  -2 -1
# FLO    2   9  13  -1    347   0  0  -6    0   0   6   7  -1 -1
# GMS    0  -1  -1   0      6   0  0   1    0   0   0  -1   0  0
# GY    89 120 347   6 157688  38 50  -3    8 -36 160 199 -40  8
# HIS    1   3   0   0     38   8  4  -5    0  -2   4   6  -2  0
# HW     1   1   0   0     50   4  4  -1    0  -1   3   4   0  0
# NGS   -2  -7  -6   1     -3  -5 -1  11    0   2  -5  -7   2  1
# NGSP   0   0   0   0      8   0  0   0    0   0   0   0   0  0
# NSS    0  -1   0   0    -36  -2 -1   2    0   1  -1  -2   1  0
# PH     0  14   6   0    160   4  3  -5    0  -1  17  18  -2 -1
# SH     0  16   7  -1    199   6  4  -7    0  -2  18  20  -2 -1
# SL     0  -2  -1   0    -40  -2  0   2    0   1  -2  -2   0  0
# SW     0  -1  -1   0      8   0  0   1    0   0  -1  -1   0  0

pcov <- gmd(mod, what = "pcov")
# Class of the model: gamem
# Variable extracted: pcov
round_cols(pcov, digits = 0) # to fit in the page width
#      FLO PH  SH FLH DIS     GY HW NSS NGSP  SL  SW  NGS GMS HIS
# FLO   14  6   7  10   2    380  0   0    0  -1  -1   -6  -1   0
# PH     6 22  23  18   0     61  3  -1    0  -1  -1   -4   0   4
# SH     7 23  25  20   1    106  4  -2    0  -2  -1   -6  -1   6
# FLH   10 18  20  20   1     56  1  -1    0  -2  -1   -7  -1   3
```

```
# DIS     2  0   1   1   1      77  1   0    0   0  0   -2   0   1
# GY    380 61 106  56  77 219250 49 -35    1 -46  1 -100   0   8
# HW      0  3   4   1   1      49  6  -1    0  -1  0   -2   0   4
# NSS     0 -1  -2  -1   0     -35 -1   1    0   1  0    3   0  -2
# NGSP    0  0   0   0   0       1  0   0    0   0  0    1   0   0
# SL     -1 -1  -2  -2   0     -46 -1   1    0   1  0    2   0  -2
# SW     -1 -1  -1  -1   0       1  0   0    0   0  0    1   0   0
# NGS    -6 -4  -6  -7  -2    -100 -2   3    1   2  1   17   1  -4
# GMS    -1  0  -1  -1   0       0  0   0    0   0  0    1   0   0
# HIS     0  4   6   3   1       8  4  -2    0  -2  0   -4   0  11
```

###### 1.4.1.4 Phenotypic and genotypic variance-covariance matrices

###### 1.4.1.5 MGIDI

1. Mannualy rescale the matrix of BLUPs

```
blups <- gmd(mod, "blupg")
# Class of the model: gamem
# Variable extracted: blupg
blups  %>% select(HIS) # original values
# # A tibble: 44 x 1
#      HIS
#    <dbl>
#  1  74.6
#  2  72.1
#  3  70.9
#  4  76.9
#  5  77.9
#  6  78.6
#  7  74.9
#  8  73.7
#  9  70.1
# 10  76.7
# # ... with 34 more rows

# The first four traits are desired to decrease.
# The Last 10 ones are desired to increase.
res <- c(rep(0, 4), rep(100, 10))
for (i in 2:ncol(blups)) {
  blups[i] <-
    resca(values = blups[i], new_max = res[i - 1], new_min = 100 - res[i - 1])
}
```

```
# rescaled values
blups
# # A tibble: 44 x 15
#    GEN     FLO    PH    SH   FLH   DIS    GY    HW   NSS  NGSP    SL    SW   NGS
#    <chr> <dbl> <dbl> <dbl> <dbl> <dbl> <dbl> <dbl> <dbl> <dbl> <dbl> <dbl> <dbl>
#  1 G1     17.5 35.1  35.5   20.2  66.7  24.2  58.9 31.7  62.4   32.6 23.8   49.7
#  2 G10    52.5 93.6  93.4   83.7  75    43.2  30.9 78.9   5.84  52.6 55.4   41.6
#  3 G11    52.5 61.4  66.1   62.6  83.3  88.1  55.5 84.5   4.74  70.5 69.9   44.2
#  4 G12    90.  65.8  71.4   68.9  58.3  58.5  73.6 67.7  31.0   77.9 90.9   56.6
#  5 G13    90.   1.96  1.66  14.6  66.7  47.7  71.2  6.21 78.1   13.7  5.06   0
#  6 G14   100.  40.7  39.7   58.4  33.3  23.7  71.3 16.1  36.5   25.3 64.3   21.7
#  7 G15    45.  41.6  45.7   55.7  58.3  60.4  47.2 77.0  40.9   57.9 61.3   74.0
#  8 G16    90.  75.3  73.9   75.2  25    59.3  45.9 50.9  47.4   37.9 58.0   56.4
#  9 G17    65.  71.4  79.9   65.1  58.3  27.9  46.8 90.1  51.5   98.9 71.1   93.9
# 10 G18    92.5 40.6  45.2   56.0  50    68.9  72.2 40.4  69.0   58.9 81.0   67.1
# # ... with 34 more rows, and 2 more variables: GMS <dbl>, HIS <dbl>
```

2. The MGIDI index

```
mgidi_index <-
  mgidi(mod,
        ideotype = c(rep("l", 4), rep("h", 10)),
        SI = 15)
#
# -------------------------------------------------------------------------------
# Principal Component Analysis
# -------------------------------------------------------------------------------
# # A tibble: 14 x 4
#    PC    Eigenvalues `Variance (%)` `Cum. variance (%)`
#    <chr>       <dbl>          <dbl>                <dbl>
#  1 PC1          5.57          39.8                  39.8
#  2 PC2          2.41          17.2                  57.0
#  3 PC3          1.82          13.0                  70.0
#  4 PC4          1.38           9.83                 79.8
#  5 PC5          1.01           7.19                 87.0
#  6 PC6          0.7           5                     92.0
#  7 PC7          0.44           3.12                 95.1
#  8 PC8          0.32           2.31                 97.4
#  9 PC9          0.16           1.16                 98.6
# 10 PC10         0.11          0.77                  99.4
# 11 PC11         0.07          0.49                  99.9
# 12 PC12         0.02          0.13                 100
# 13 PC13         0             0                    100
```

```
# 14 PC14         0              0                  100
# ---------------------------------------------------------------------------------
# Factor Analysis - factorial loadings after rotation-
# ---------------------------------------------------------------------------------
# # A tibble: 14 x 8
#    VAR      FA1   FA2    FA3   FA4    FA5 Communality Uniquenesses
#    <chr>  <dbl> <dbl>  <dbl> <dbl>  <dbl>       <dbl>        <dbl>
#  1 FLO    -0.12  0.01  0.4    0.72  0.35         0.83         0.17
#  2 PH     -0.08 -0.17  0.97   0.01 -0.01         0.97         0.03
#  3 SH     -0.21 -0.19  0.95   0.03  0.01         0.98         0.02
#  4 FLH    -0.26  0.07  0.9    0.24  0.03         0.95         0.05
#  5 DIS     0.14  0.56  0.03  -0.62 -0.290        0.81         0.19
#  6 GY      0.03  0     0     -0.01 -0.93         0.87         0.13
#  7 HW      0.08  0.85 -0.18   0.05  0.04         0.76         0.24
#  8 NSS    -0.88 -0.26  0.06  -0.2   0.07         0.88         0.12
#  9 NGSP    0.04  0.14 -0.01   0.87 -0.17         0.8          0.2
# 10 SL     -0.84 -0.2   0.290  0.12  0.11         0.87         0.13
# 11 SW     -0.71  0.26  0.39   0.48 -0.03         0.95         0.05
# 12 NGS    -0.71 -0.1   0.09   0.54 -0.09         0.83         0.17
# 13 GMS    -0.580  0.42  0.36   0.54 -0.01         0.94         0.06
# 14 HIS     0.54  0.65 -0.16   0.17  0.02         0.77         0.23
# ---------------------------------------------------------------------------------
# Comunalit Mean: 0.8700916
# ---------------------------------------------------------------------------------
# Selection differential
# ---------------------------------------------------------------------------------
# # A tibble: 14 x 11
#    VAR   Factor     Xo     Xs       SD  SDperc    h2       SG  SGperc sense
#    <chr> <chr>   <dbl>  <dbl>    <dbl>   <dbl> <dbl>    <dbl>   <dbl> <chr>
#  1 NSS   FA 1   1.55e1 1.60e1   0.450    2.90  0.765   0.345    2.22  incr~
#  2 SL    FA 1   8.63e0 9.12e0   0.498    5.78  0.668   0.333    3.86  incr~
#  3 SW    FA 1   2.17e0 2.54e0   0.371   17.1   0.836   0.310   14.3   incr~
#  4 NGS   FA 1   4.05e1 4.20e1   1.51     3.73  0.655   0.990    2.45  incr~
#  5 GMS   FA 1   1.62e0 1.88e0   0.259   16.0   0.812   0.210   13.0   incr~
#  6 HW    FA 2   7.63e1 7.77e1   1.37     1.79  0.801   1.10     1.44  incr~
#  7 HIS   FA 2   7.47e1 7.42e1  -0.523   -0.700 0.788  -0.412   -0.551 incr~
#  8 PH    FA 3   8.65e1 8.64e1  -0.0756  -0.0875 0.784 -0.0593  -0.0686 decr~
#  9 SH    FA 3   7.78e1 7.71e1  -0.698   -0.897 0.821  -0.573   -0.736 decr~
# 10 FLH   FA 3   6.04e1 5.87e1  -1.68    -2.78  0.826  -1.39    -2.29  decr~
# 11 FLO   FA 4   6.05e1 5.91e1  -1.34    -2.22  0.923  -1.24    -2.05  decr~
# 12 DIS   FA 4   2.84e0 3.11e0   0.271    9.53  0.896   0.243    8.54  incr~
# 13 NGSP  FA 4   2.62e0 2.64e0   0.0235   0.896 0.516   0.0121   0.462 incr~
# 14 GY    FA 5   4.38e3 4.62e3 241.       5.50  0.719 173.       3.96  incr~
# # ... with 1 more variable: goal <dbl>
```

```
# -----------------------------------------------------------------------------
# Selected genotypes
# -----------------------------------------------------------------------------
# G12 G37 G18 G8 G11 G9 G32
# -----------------------------------------------------------------------------
```

```
# MGIDI index
plot(mgidi_index)
```

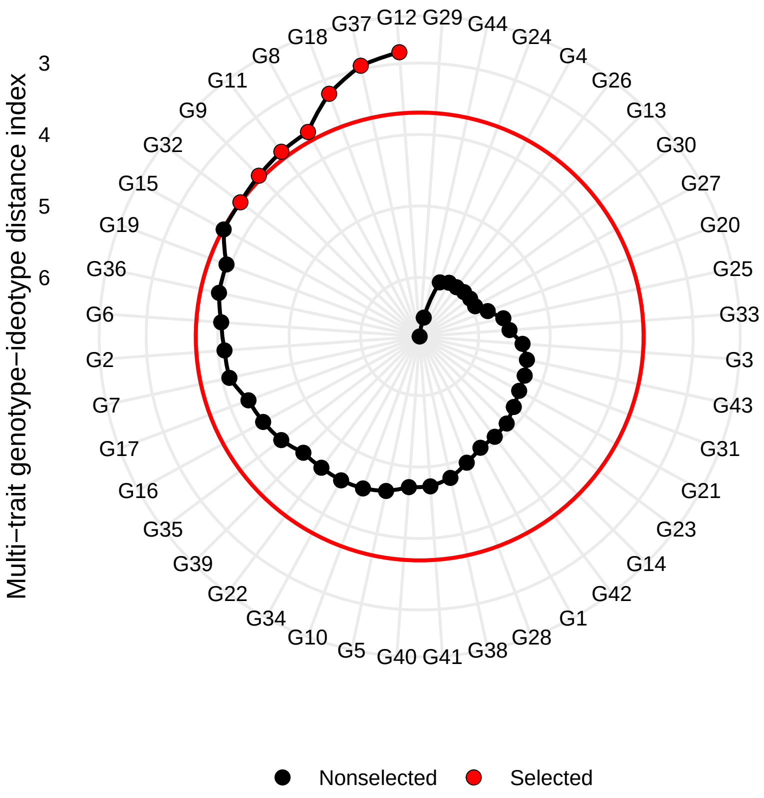

```
# Contribution (selected genotypes)
plot(mgidi_index,
     type = "contribution",
     width.bar = 1,
     size.line = 0.2)
```

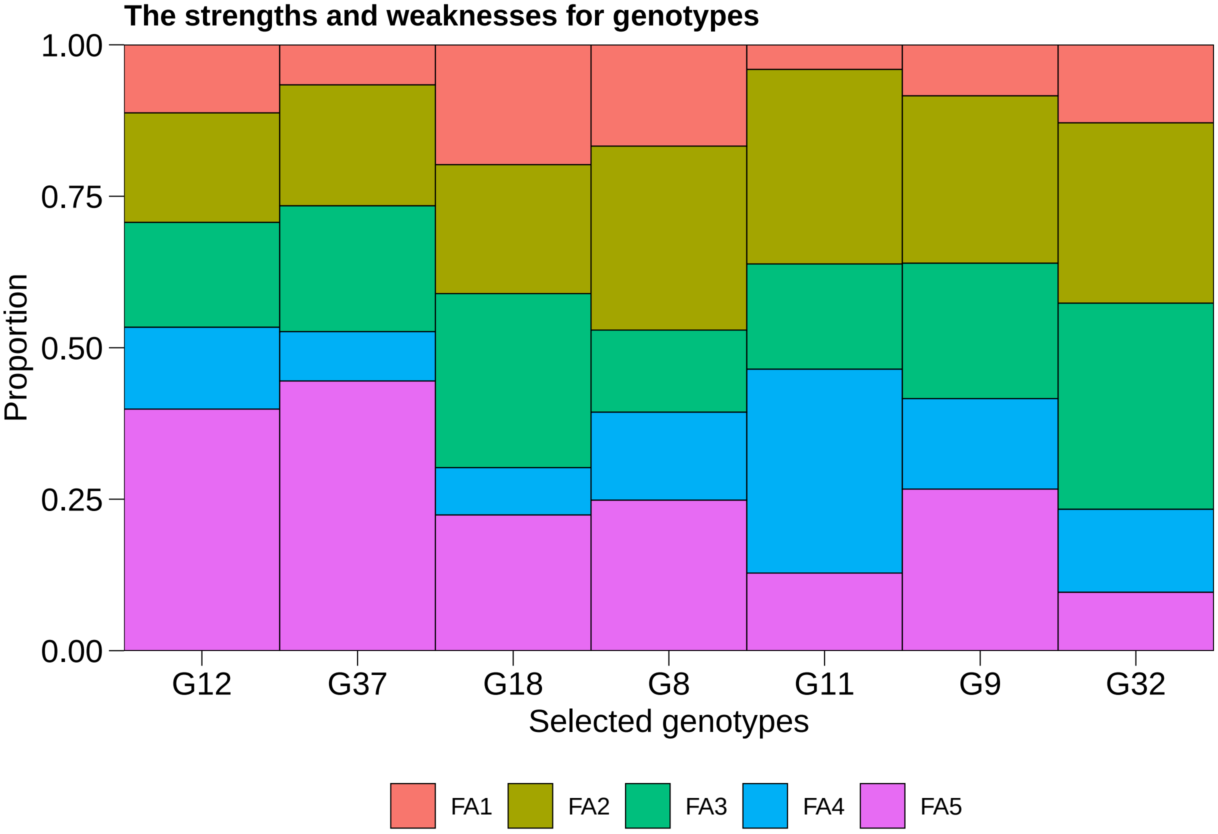

**1.4.1.6 FAI-BLUP** For the FAI-BLUP index (Rocha, Machado, & Carneiro, 2018), the desired ideotype (DI) was defined by a vector with "min" for the traits in which lower values are desired and "max" for the traits in which higher values are desired. Then the distances from each genotype according to ideotypes were estimated and converted into spatial probability as follows:

$$P_{ij} = \frac{\frac{1}{d_{ij}}}{\sum\limits_{i=1;j=1}^{i=n;j=m} \frac{1}{d_{ij}}} \quad (1)$$

where $P_{ij}$ is the probability of the $i$th genotype ($i = 1, 2, ..., n$) to be similar to the $j$th ideotype ($j = 1, 2, ..., m$); $d_{ij}$ is the genotype-ideotype distance from the $i$th genotype to the $j$th ideotype based on standardized mean Euclidean distance.

```
fai <-
  fai_blup(mod,
           DI = c(rep("min", 4), rep("max", 10)),
           UI = c(rep("max", 4), rep("min", 10)),
           SI = 15)
#
# -----------------------------------------------------------------------------
```

```
# Principal Component Analysis
# -----------------------------------------------------------------------------------
#      eigen.values cumulative.var
# PC1          5.57          39.76
# PC2          2.41          56.99
# PC3          1.82          69.99
# PC4          1.38          79.82
# PC5          1.01          87.01
# PC6          0.70          92.01
# PC7          0.44          95.14
# PC8          0.32          97.44
# PC9          0.16          98.61
# PC10         0.11          99.38
# PC11         0.07          99.87
# PC12         0.02         100.00
# PC13         0.00         100.00
# PC14         0.00         100.00
#
# -----------------------------------------------------------------------------------
# Factor Analysis
# -----------------------------------------------------------------------------------
#        FA1   FA2   FA3   FA4   FA5 comunalits
# FLO  -0.12  0.01 -0.40 -0.72 -0.35       0.83
# PH   -0.08 -0.17 -0.97 -0.01  0.01       0.97
# SH   -0.21 -0.19 -0.95 -0.03 -0.01       0.98
# FLH  -0.26  0.07 -0.90 -0.24 -0.03       0.95
# DIS  -0.14 -0.56  0.03 -0.62 -0.29       0.81
# GY   -0.03  0.00  0.00 -0.01 -0.93       0.87
# HW   -0.08 -0.85 -0.18  0.05  0.04       0.76
# NSS   0.88  0.26  0.06 -0.20  0.07       0.88
# NGSP -0.04 -0.14 -0.01  0.87 -0.17       0.80
# SL    0.84  0.20  0.29  0.12  0.11       0.87
# SW    0.71 -0.26  0.39  0.48 -0.03       0.95
# NGS   0.71  0.10  0.09  0.54 -0.09       0.83
# GMS   0.58 -0.42  0.36  0.54 -0.01       0.94
# HIS  -0.54 -0.65 -0.16  0.17  0.02       0.77
#
# -----------------------------------------------------------------------------------
# Comunalit Mean: 0.8700916
# Selection differential
# -----------------------------------------------------------------------------------
# # A tibble: 14 x 11
#    VAR   Factor      Xo     Xs       SD SDperc    h2       SG SGperc sense  goal
#    <chr>  <dbl>   <dbl>  <dbl>    <dbl>  <dbl> <dbl>    <dbl>  <dbl> <chr> <dbl>
```

```
#  1 NSS        1   15.5  1.55e1  -0.0271 -0.174 0.765  -0.0207 -0.133 incr~     0
#  2 SL         1    8.63 8.78e0   0.158   1.83  0.668   0.106   1.22  incr~   100
#  3 SW         1    2.17 2.50e0   0.332  15.3   0.836   0.278  12.8   incr~   100
#  4 NGS        1   40.5  4.30e1   2.49    6.16  0.655   1.63    4.04  incr~   100
#  5 GMS        1    1.62 1.89e0   0.271  16.8   0.812   0.220  13.6   incr~   100
#  6 HW         2   76.3  7.73e1   1.02    1.33  0.801   0.815   1.07  incr~   100
#  7 HIS        2   74.7  7.58e1   1.10    1.47  0.788   0.866   1.16  incr~   100
#  8 PH         3   86.5  8.61e1  -0.367  -0.424 0.784  -0.287  -0.332 decr~   100
#  9 SH         3   77.8  7.72e1  -0.585  -0.751 0.821  -0.480  -0.616 decr~   100
# 10 FLH        3   60.4  5.83e1  -2.08   -3.45  0.826  -1.72   -2.85  decr~   100
# 11 FLO        4   60.5  5.78e1  -2.66   -4.40  0.923  -2.46   -4.07  decr~   100
# 12 DIS        4    2.84 2.77e0  -0.0708 -2.49  0.896  -0.0635 -2.23  incr~     0
# 13 NGSP       4    2.62 2.74e0   0.122   4.68  0.516   0.0631  2.41  incr~   100
# 14 GY         5 4380.   4.62e3 238.      5.43  0.719 171.      3.91  incr~   100
#
# ---------------------------------------------------------------------------------
# Selected genotypes
# G37 G18 G12 G32 G7 G6 G19
# ---------------------------------------------------------------------------------
```

**1.4.1.7 Smith-Hazel** The SH index was computed according to (2)

$$\mathbf{b} = \mathbf{P}^{-1}\mathbf{G}\mathbf{w} \tag{2}$$

where **P** and **G** are phenotypic and genetic variance-covariance matrices, respectively, and **b** and **w** are vectors of index coefficients and economic weightings, respectively. All traits were considered as having the same economic weight ($\mathbf{w} = 1$) with negative sign (-1) for traits in which lower values are desired and positive sign (1) for traits in which higher values are desired, thus, enabling to place all traits data in the desired direction for selection (Smiderle et al., 2019).

The genetic worth (**I**) of the individual genotype based on the evaluated traits were then computed as follows:

$$\mathbf{I} = \mathbf{X}\mathbf{b} \tag{3}$$

where **X** is an $g \times p$ matrix of BLUPs for the $g$ genotypes and $p$ traits and **b** is a $1 \times p$ vector of index coefficient for the $p$ traits.

Two scenarios were considered. In the first one (SH-1), the SH index was computed with all the 14 traits. In the second one (SH-2), the multicollinearity-generating traits were excluded and the index was computed with the remaining traits. The traits to be excluded were chosen with the function `non_collinear_vars()`

1. All variables

```r
smith_index <-
  Smith_Hazel(mod,
              weights = c(rep(-1, 4), rep(1, 10)),
              SI = 15)
```

2. Remove colinear variables

```r
non_collinear_vars(data)
#          Parameter                                                    values
# 1       Predictors                                                        12
# 2              VIF                                                     9.647
# 3 Condition Number                                                    75.973
# 4      Determinant                                              0.0004347741
# 5         Selected GY, HW, DIS, HIS, SL, FLO, SW, NGSP, PH, NSS, FLH, NGS
# 6          Removed                                                   SH, GMS

# create a new data set
df_colin <- remove_cols(data, SH, GMS)
mod_colin <- gamem(df_colin,
                   gen = GEN,
                   rep = BLOCK,
                   resp = everything())
# Method: REML/BLUP
# Random effects: GEN
# Fixed effects: REP
# Denominador DF: Satterthwaite's method
# ---------------------------------------------------------------------------
# P-values for Likelihood Ratio Test of the analyzed traits
# ---------------------------------------------------------------------------
#     model      FLO        PH       FLH      DIS        GY        HW       NSS
#  Complete       NA        NA        NA       NA        NA        NA        NA
#  Genotype 5.31e-23 1.66e-09 6.21e-12 1.36e-18 5.71e-07 2.19e-10 1.15e-08
#     NGSP       SL       SW       NGS      HIS
#       NA       NA       NA        NA       NA
#  0.00465 1.45e-05 1.3e-12 2.89e-05 1.08e-09
# ---------------------------------------------------------------------------
# All variables with significant (p < 0.05) genotype effect
```

3. Index after removing colinear variables

```r
smith_index_colin <-
  Smith_Hazel(mod_colin,
              weights = c(rep(-1, 3), rep(1, 9)),
              SI = 15)
```

```
sg_mgidi <-
  mgidi_index$sel_dif %>%
  select_cols(Factor, VAR, Xo, sense, h2, SGperc) %>%
  rename(MGIDI = SGperc)
sg_fai <-
  fai$selection_diferential$ID1 %>%
  select_cols(VAR, SGperc) %>%
  rename(FAI_BLUP = SGperc)
sg_hz_all <-
  smith_index$sel_dif %>%
  select_cols(VAR, SGperc) %>%
  rename(HZ1 = SGperc)
sg_hz_colin <-
  smith_index_colin$sel_dif %>%
  select_cols(VAR, SGperc) %>%
  rename(HZ2 = SGperc)

# BLUPs
blups <-
  gmd(mod, "blupg") %>%
  desc_stat(stats = c("mean, se")) %>%
  round_cols() %>%
  add_cols(mse = paste(mean, "+-", se, sep = "")) %>%
  select(VAR = variable, mse = mse)
# Class of the model: gamem
# Variable extracted: blupg

# join all selection indexes
selection_gain <-
  left_join(sg_mgidi, blups, by = "VAR") %>%
  left_join(sg_fai, by = "VAR") %>%
  left_join(sg_hz_all, by = "VAR") %>%
  left_join(sg_hz_colin, by = "VAR") %>%
  select_cols(Factor, VAR, mse, MGIDI, FAI_BLUP, HZ1, HZ2) %>%
  tidy_strings(Factor, sep = "")

selection_gain
# # A tibble: 14 x 7
#    Factor VAR   mse                MGIDI FAI_BLUP    HZ1     HZ2
#    <chr>  <chr> <chr>              <dbl>    <dbl>  <dbl>   <dbl>
```

```
#   1 FA1    NSS   15.54+-0.14     2.22      -0.133 -4.05  -1.87
#   2 FA1    SL    8.63+-0.08      3.86       1.22  -4.71  -2.20
#   3 FA1    SW    2.17+-0.04     14.3       12.8   -8.21   3.40
#   4 FA1    NGS   40.47+-0.41     2.45       4.04  -3.16   0.997
#   5 FA1    GMS   1.62+-0.03     13.0       13.6   -6.24  NA
#   6 FA2    HW    76.31+-0.29     1.44       1.07   1.11   0.348
#   7 FA2    HIS   74.71+-0.39    -0.551      1.16   1.71   1.08
#   8 FA3    PH    86.46+-0.55    -0.0686    -0.332  3.86   0.104
#   9 FA3    SH    77.83+-0.62    -0.736     -0.616  5.49  NA
#  10 FA3    FLH   60.42+-0.56    -2.29      -2.85   4.89  -0.0283
#  11 FA4    FLO   60.45+-0.52    -2.05      -4.07   4.40   1.10
#  12 FA4    DIS   2.84+-0.14      8.54      -2.23  20.7   11.2
#  13 FA4    NGSP  2.62+-0.02      0.462      2.41   0.419  1.43
#  14 FA5    GY    4379.69+-50.77  3.96       3.91   4.64   7.62
```

###### 1.4.1.8 Binding the results

##### 1.4.2 Dataset 2 - Nitrogen and Sulfur management

In this section, we extend the theory of the MGIDI index to evaluate an experiment (Olivoto et al., 2016) with a two-way qualitative-qualitative treatment structure. The focus is to choose the best treatment that allows desired values for most of the evaluated traits. Here, the goal is to obtain low value for P and PL and high values for the remaining traits. Note that the input needed in the function `mgidi()` is a two-way table with treatment/genotypes in the row names and traits in the columns.

```
# Pass column TRAT to row names
data_ns_mat <-
  column_to_rownames(data_ns, "TRAT")

# Define the ideotype vector
ide_vect <- c("l", "l", rep("h", 8))
ide_vect
#  [1] "l" "l" "h" "h" "h" "h" "h" "h" "h" "h"

# Fit the mgidi index
mgidi_data_ns <-
  mgidi(data_ns_mat,
        ideotype = ide_vect)
#
# ------------------------------------------------------------------------------------
# Principal Component Analysis
# ------------------------------------------------------------------------------------
# # A tibble: 10 x 4
```

```
#    PC    Eigenvalues `Variance (%)` `Cum. variance (%)`
#    <chr>       <dbl>          <dbl>               <dbl>
#  1 PC1          4.3           43.0                 43.0
#  2 PC2          3.03          30.3                 73.3
#  3 PC3          1.26          12.6                 85.9
#  4 PC4          0.67           6.68                92.6
#  5 PC5          0.59           5.93                98.5
#  6 PC6          0.12           1.15                99.7
#  7 PC7          0.03           0.32               100
#  8 PC8          0              0                  100
#  9 PC9          0              0                  100
# 10 PC10         0              0                  100
# ------------------------------------------------------------------------------
# Factor Analysis - factorial loadings after rotation-
# ------------------------------------------------------------------------------
# # A tibble: 10 x 6
#    VAR     FA1   FA2   FA3 Communality Uniquenesses
#    <chr> <dbl> <dbl> <dbl>       <dbl>        <dbl>
#  1 P      0.63 -0.73  0.08        0.93         0.07
#  2 PL     0.9  -0.09  0.26        0.89         0.11
#  3 L      0.88  0.18  0.3         0.9          0.1
#  4 TIL   -0.24 -0.63 -0.71        0.96         0.04
#  5 SSM    0.03 -0.16 -0.96        0.95         0.05
#  6 GY    -0.2  -0.04 -0.97        0.99         0.01
#  7 HW     0.53 -0.49 -0.34        0.63         0.37
#  8 W      0.37  0.87  0.21        0.94         0.06
#  9 GLU    0.84  0.05  0.15        0.73         0.27
# 10 PROT  -0.79 -0.06  0.19        0.67         0.33
# ------------------------------------------------------------------------------
# Comunalit Mean: 0.8591681
# ------------------------------------------------------------------------------
# Selection differential
# ------------------------------------------------------------------------------
# # A tibble: 10 x 8
#    VAR   Factor     Xo     Xs       SD  SDperc sense     goal
#    <chr> <chr>   <dbl>  <dbl>    <dbl>   <dbl> <chr>    <dbl>
#  1 PL    FA 1     1.21   1.14 -0.0663 -5.49   decrease   100
#  2 L     FA 1    81.7   85.2   3.59    4.40   increase   100
#  3 HW    FA 1    79.2   79.2   0.0312  0.0394 increase   100
#  4 GLU   FA 1    28.7   29.8   1.12    3.91   increase   100
#  5 PROT  FA 1    12.3   11.9  -0.405  -3.28   increase     0
#  6 P     FA 2    97.2   96.8  -0.469  -0.482  decrease   100
#  7 W     FA 2   256.   264.    7.78    3.04   increase   100
#  8 TIL   FA 3     4.84   5.48  0.635  13.1    increase   100
```

```
#  9 SSM    FA 3     663.     794.    132.      19.9     increase   100
# 10 GY     FA 3    5941.    6459     518.       8.73    increase   100
# -------------------------------------------------------------------------------
# Selected genotypes
# --------------------------------------------------------------------------------
# WS_30T_40DR_30B
# --------------------------------------------------------------------------------

# Radar plot with the ranking for treatments
plot(mgidi_data_ns)
```

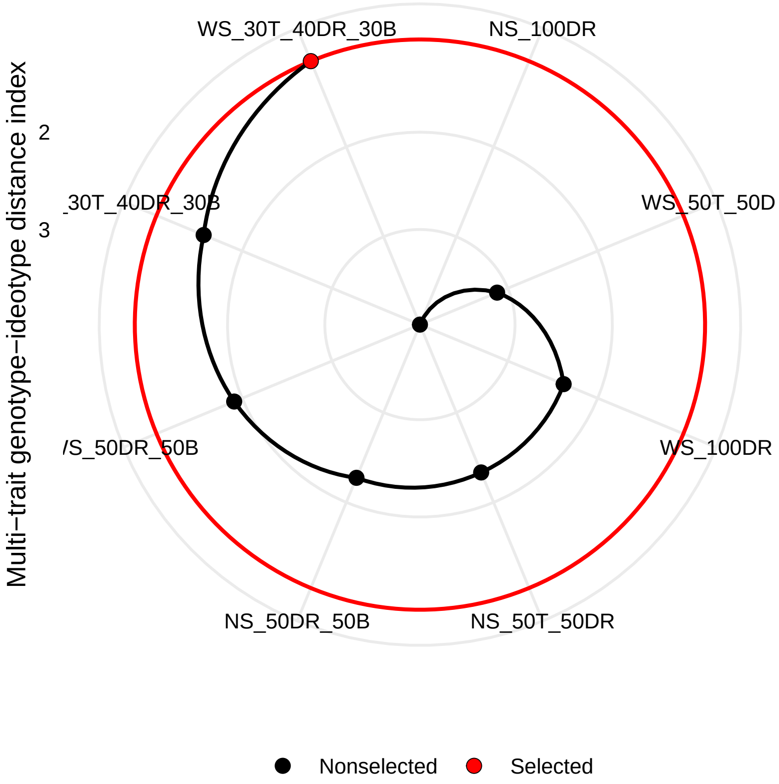

Three factors were retained, explaining 85.9% of the total variation. The traits grouped into each factor where: **FA1**: PL, L, HW, GLU, and PROT. **FA2**: P and W; and **FA3**, TIL, SSM, and GY.

The MGIDI index indicated the treatment **WS_30T_40DR_30B** (30% N at the tillering stage, 40% N at the double ring stage, and 30% N at the boosting stage with sulfur implementation) as the best performing treatment. This treatment provides desired values (lower or higher value) for 9 of the 10 analyzed traits, thus, the success rate, in this case, was 90%. The treatment ranking also suggests that nitrogen applied at once in double ring or in two applications (50% N at the tillering and 50% N at the double ring stage), independently on the sulfur management, are the poorly performing treatments.

The strengths and weaknesses view of treatments shown below indicates that the selected treatment (**WS_30T_40DR_30B**) performed well for the traits into FA1 and FA3, which can be seen in the plot showing the phenotypic means.

```
plot(mgidi_data_ns,
     type = "contribution",
     genotypes = "all",
     x.lab = "Treatments",
     width = 1,
     title = "The strengths and weaknesses view of treatments",
     rotate = TRUE)
```

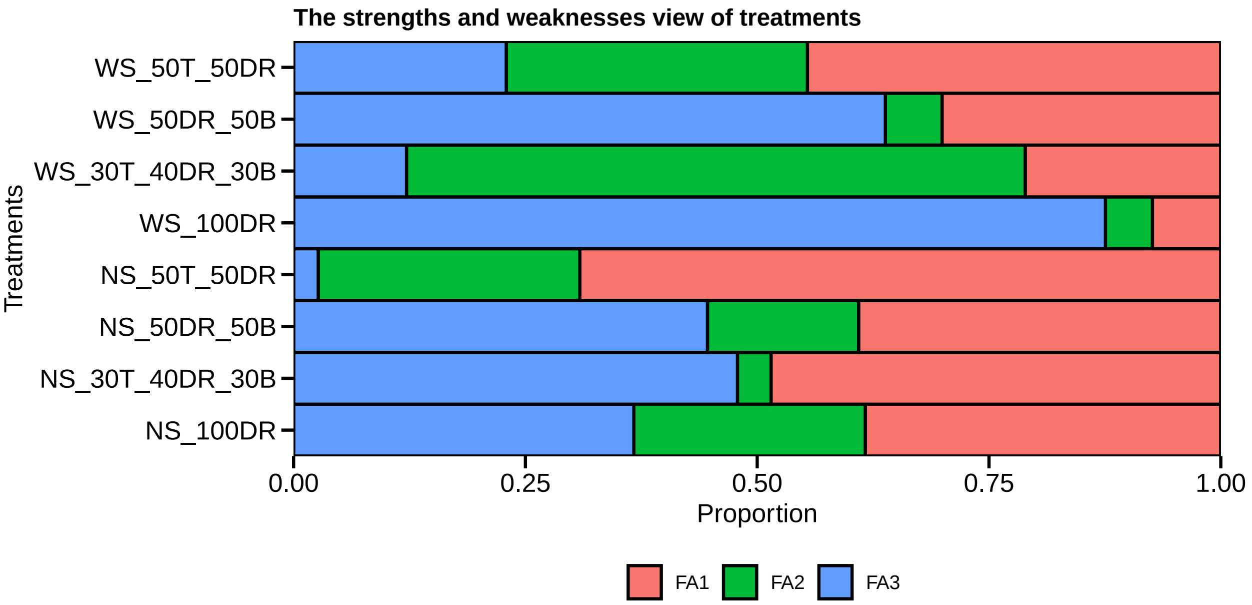

```
# Generate a plot to show the mean for each treatment
library(ggplot2)
df_plot <-
  data_ns %>%
  pivot_longer(-TRAT)

means <- means_by(df_plot, name)

ggplot(df_plot, aes(value, TRAT)) +
  geom_bar(stat = "identity",
           fill = rep(c(rep("gray", 1), "blue", rep("gray", 6)), 10),
```

```
           color = "black",
           width = 1) +
  facet_wrap(~name, scales = "free_x", ncol = 2) +
  geom_vline(data = means, aes(xintercept = value), color = "red")+
  scale_x_continuous(expand = expansion(c(0, 0.05)))
```

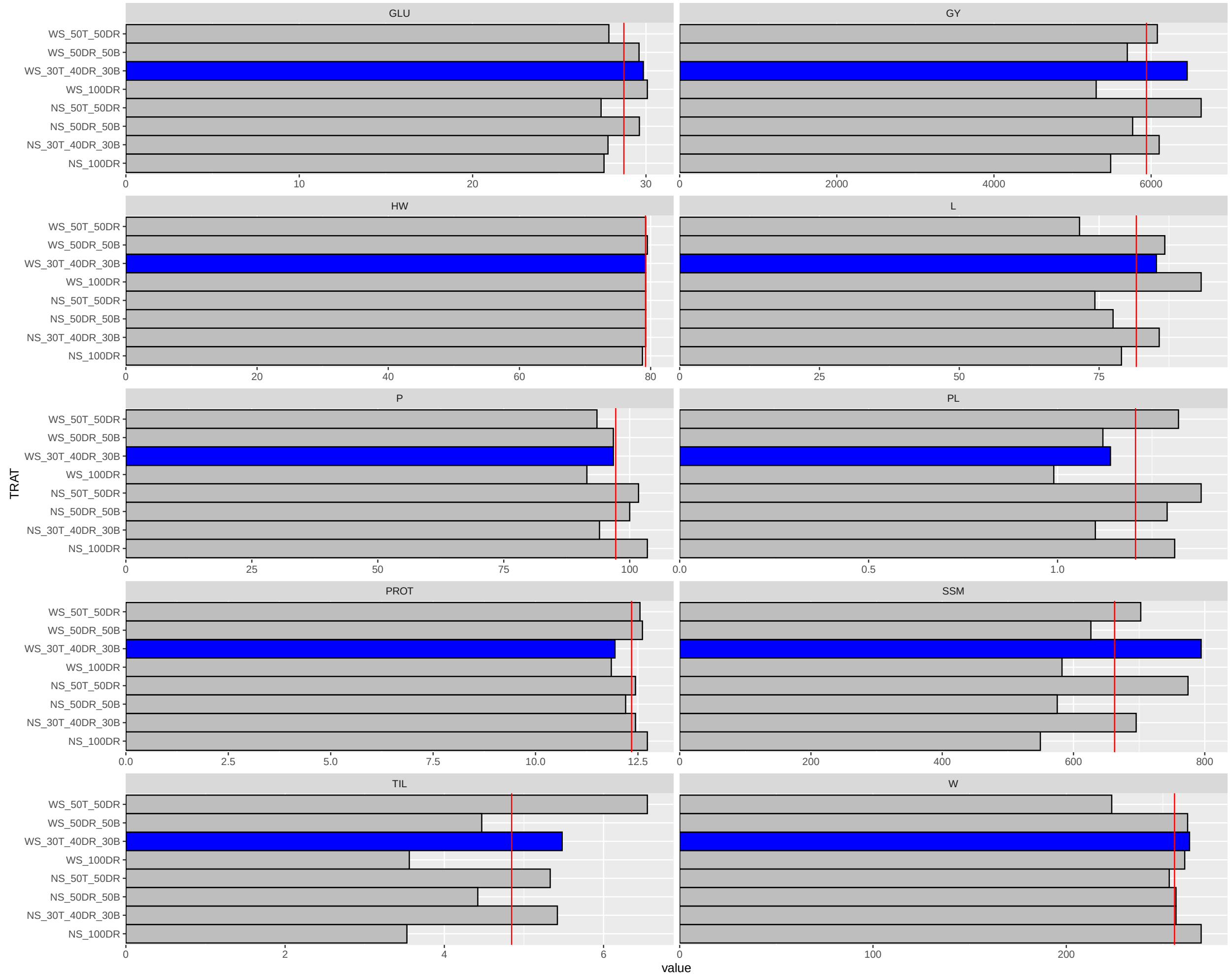

Figure S 1: Phenotypic means for each treatment the treatment selected by the MGIDI index (WS_30T_40DR_30B) is highlighted in blue.

#### 1.5 A step-by-step guide for future studies

The mgidi index can be computed in two main ways: Using a two-way table, e.g, genotype-by-trait or treatment-by trait means or a raw data. The function `mgidi()` has the following arguments.

```
args(mgidi)
# function (.data, use_data = "blup", SI = 15, mineval = 1, ideotype = NULL,
#     use = "complete.obs", verbose = TRUE)
# NULL
```

- `.data`:An object fitted with the function `gamem()` (mixed-effect model), `gafem()` (fixed effect model), or a two-way table with genotypes/treatments in rows and trait in columns. Please, ensure that genotypes/treatments labels are given as row names.
- `use_data`: Define which data to use If `.data` is an object of class`gamem`. Defaults to `"blup"` (the BLUPs are used). Use `"pheno"` to use phenotypic means instead BLUPs for computing the index.
- `SI`: An integer (0-100). The selection intensity in percentage of the total number of genotypes/treatments.
- `mineval`: The minimum value so that an eigenvalue is retained in the factor analysis. Defaults to 1.
- `ideotype`: A vector with length equal to the number of variables used to plan the ideotype. Use `"h"` to indicate the traits in which higher values are desired or `"l"` to indicate the variables in which lower values are desired. For example, `ideotype = c("h, h, h, h, l")` will consider that the ideotype has higher values for the first four traits and lower values for the last trait. It is also possible to use `"m"` to define an ideotype in which mean values are desirable. If `.data` is a model fitted with the functions `gafem()` or `gamem()`, the order of the traits will be declared in the argument `resp` in those functions.
- `use`: The method for computing (CO)variances in the presence of missing values. Defaults to `"complete.obs"`, i.e., missing values are handled by casewise deletion.
- `verbose`: If `verbose = TRUE` (Default) then some results are shown in the console.

In our real Dataset 1 example, we defined the ideotype as a vector with four `"h"`s and ten `"l"`s.

```
ideotype_vector <- c(rep("l", 4), rep("h", 10))
ideotype_vector
#  [1] "l" "l" "l" "l" "h" "h" "h" "h" "h" "h" "h" "h" "h" "h"
```

> ***Quick tip!*** *we may reorder the columns with `reorder_cols()` to put columns with similar desired gains together to avoid mixing `"l"`s and `"h"`s in the ideotype vector.*

##### 1.5.1 Using a two-way table with trait means

In this example, we will create a two-way table using the raw data 'data' that will contains the phenotypic means.

```
data_mean <-
  means_by(data, GEN) %>%
  column_to_rownames("GEN")

data_mean %>% round_cols(digits = 1) %>% head()
#      FLO   PH   SH  FLH DIS     GY   HW  NSS NGSP  SL  SW  NGS GMS  HIS
# G1  67.0 90.1 82.0 66.6 3.7 3921.7 76.8 14.6  2.8 8.1 1.9 40.2 1.4 74.6
# G10 62.3 79.2 70.4 54.9 4.0 4287.7 74.2 17.1  2.3 8.8 2.3 38.9 1.6 71.4
# G11 62.3 85.2 75.9 58.8 4.3 5152.3 76.5 17.4  2.3 9.3 2.5 39.3 1.7 69.8
# G12 57.3 84.4 74.8 57.6 3.3 4582.6 78.2 16.5  2.5 9.6 2.8 41.4 2.1 77.5
# G13 57.3 96.3 88.8 67.6 3.7 4373.9 78.0 13.2  2.9 7.5 1.6 31.8 1.3 78.8
# G14 56.0 89.1 81.2 59.6 2.3 3912.6 78.0 13.7  2.6 7.9 2.4 35.5 1.9 79.7
```

Now, the mgidi index can be computed with `mgidi()` using the above two-way table.

```
mgidi_mean <-
  mgidi(data_mean, # input data. In this case, a two-way table
        ideotype = ideotype_vector, # The vector planning the ideotype
        SI = 15, # The selection intensity
        verbose = T) # Results to the console?
#
# -------------------------------------------------------------------------------
# Principal Component Analysis
# -------------------------------------------------------------------------------
# # A tibble: 14 x 4
#    PC    Eigenvalues `Variance (%)` `Cum. variance (%)`
#    <chr>       <dbl>          <dbl>               <dbl>
#  1 PC1          5.57          39.8                 39.8
#  2 PC2          2.41          17.2                 57.0
#  3 PC3          1.82          13.0                 70.0
#  4 PC4          1.38           9.83                79.8
#  5 PC5          1.01           7.19                87.0
#  6 PC6          0.7            5                   92.0
#  7 PC7          0.44           3.12                95.1
#  8 PC8          0.32           2.31                97.4
#  9 PC9          0.16           1.16                98.6
# 10 PC10         0.11           0.77                99.4
# 11 PC11         0.07           0.49                99.9
# 12 PC12         0.02           0.13               100
# 13 PC13         0              0                  100
```

```
# 14 PC14         0              0                  100
# ---------------------------------------------------------------------------------------
# Factor Analysis - factorial loadings after rotation-
# ---------------------------------------------------------------------------------------
# # A tibble: 14 x 8
#    VAR      FA1   FA2    FA3   FA4    FA5 Communality Uniquenesses
#    <chr>  <dbl> <dbl>  <dbl> <dbl>  <dbl>       <dbl>        <dbl>
#  1 FLO    -0.12  0.01  0.4    0.72  0.35         0.83         0.17
#  2 PH     -0.08 -0.17  0.97   0.01 -0.01         0.97         0.03
#  3 SH     -0.21 -0.19  0.95   0.03  0.01         0.98         0.02
#  4 FLH    -0.26  0.07  0.9    0.24  0.03         0.95         0.05
#  5 DIS     0.14  0.56  0.03  -0.62 -0.290        0.81         0.19
#  6 GY      0.03  0     0     -0.01 -0.93         0.87         0.13
#  7 HW      0.08  0.85 -0.18   0.05  0.04         0.76         0.24
#  8 NSS    -0.88 -0.26  0.06  -0.2   0.07         0.88         0.12
#  9 NGSP    0.04  0.14 -0.01   0.87 -0.17         0.8          0.2
# 10 SL     -0.84 -0.2   0.290  0.12  0.11         0.87         0.13
# 11 SW     -0.71  0.26  0.39   0.48 -0.03         0.95         0.05
# 12 NGS    -0.71 -0.1   0.09   0.54 -0.09         0.83         0.17
# 13 GMS    -0.580  0.42  0.36   0.54 -0.01         0.94         0.06
# 14 HIS     0.54  0.65 -0.16   0.17  0.02         0.77         0.23
# ---------------------------------------------------------------------------------------
# Comunalit Mean: 0.8700916
# ---------------------------------------------------------------------------------------
# Selection differential
# ---------------------------------------------------------------------------------------
# # A tibble: 14 x 8
#    VAR   Factor     Xo      Xs       SD  SDperc sense     goal
#    <chr> <chr>   <dbl>   <dbl>    <dbl>   <dbl> <chr>    <dbl>
#  1 NSS   FA 1     15.5    16.1    0.588    3.79 increase   100
#  2 SL    FA 1      8.63    9.37   0.746    8.65 increase   100
#  3 SW    FA 1      2.17    2.61   0.444   20.5  increase   100
#  4 NGS   FA 1     40.5    42.8    2.31     5.70 increase   100
#  5 GMS   FA 1      1.62    1.94   0.319   19.7  increase   100
#  6 HW    FA 2     76.3    78.0    1.71     2.24 increase   100
#  7 HIS   FA 2     74.7    74.0   -0.664   -0.888 increase     0
#  8 PH    FA 3     86.5    86.4   -0.0965  -0.112 decrease   100
#  9 SH    FA 3     77.8    77.0   -0.850   -1.09  decrease   100
# 10 FLH   FA 3     60.4    58.4   -2.03    -3.36  decrease   100
# 11 FLO   FA 4     60.5    59     -1.45    -2.41  decrease   100
# 12 DIS   FA 4      2.84    3.14   0.302   10.6   increase   100
# 13 NGSP  FA 4      2.62    2.66   0.0455   1.74  increase   100
# 14 GY    FA 5   4380.   4715.   335.       7.65  increase   100
# ---------------------------------------------------------------------------------------
```

```
# Selected genotypes
# ---------------------------------------------------------------------------------
# G12 G37 G18 G8 G11 G9 G32
# ---------------------------------------------------------------------------------

mgidi_mean$sel_dif %>% round_cols()
# # A tibble: 14 x 8
#    VAR   Factor      Xo      Xs     SD SDperc sense     goal
#    <chr> <chr>    <dbl>   <dbl>  <dbl>  <dbl> <chr>    <dbl>
#  1 NSS   FA 1      15.5    16.1   0.59   3.79 increase   100
#  2 SL    FA 1       8.63    9.37  0.75   8.65 increase   100
#  3 SW    FA 1       2.17    2.61  0.44  20.5  increase   100
#  4 NGS   FA 1      40.5    42.8   2.31   5.7  increase   100
#  5 GMS   FA 1       1.62    1.94  0.32  19.7  increase   100
#  6 HW    FA 2      76.3    78.0   1.71   2.24 increase   100
#  7 HIS   FA 2      74.7    74.0  -0.66  -0.89 increase     0
#  8 PH    FA 3      86.5    86.4  -0.1   -0.11 decrease   100
#  9 SH    FA 3      77.8    77.0  -0.85  -1.09 decrease   100
# 10 FLH   FA 3      60.4    58.4  -2.03  -3.36 decrease   100
# 11 FLO   FA 4      60.4    59    -1.45  -2.41 decrease   100
# 12 DIS   FA 4       2.84    3.14  0.3   10.6  increase   100
# 13 NGSP  FA 4       2.62    2.66  0.05   1.74 increase   100
# 14 GY    FA 5    4380.   4715.  335.     7.65 increase   100
```

##### 1.5.2 Using the raw data

The model fitted with both `gafem()`and `gamem()` can be then be used as input data in the `mgidi()` function to compute the mgidi index. In the following example, the selection differential are the same as those observed when using the two-way table as input data. The only difference here is that the selection gains are computed because the heritability for each trait is extracted from the fitted model.

```
# Fixed-effect model for all variables in the data set
mod_fix <- gafem(data,
                 gen = GEN,
                 rep = BLOCK,
                 resp = everything(),
                 verbose = FALSE)

# Compute the mgidi index
mgidi_fix <-
  mgidi(mod_fix, # input data. In this case, the fitted model
        ideotype = ideotype_vector, # The vector planning the ideotype
        SI = 15, # The selection intensity
```

```
       verbose = FALSE) # Results to the console?

# extract the selection gains
mgidi_fix$sel_dif %>% round_cols()
# # A tibble: 14 x 11
#    VAR   Factor     Xo      Xs      SD SDperc    h2      SG SGperc sense     goal
#    <chr> <chr>   <dbl>   <dbl>   <dbl>  <dbl> <dbl>   <dbl>  <dbl> <chr>    <dbl>
#  1 NSS   FA 1     15.5    16.1    0.59   3.79  0.77    0.45   2.9  increase   100
#  2 SL    FA 1      8.63    9.37   0.75   8.65  0.67    0.5    5.78 increase   100
#  3 SW    FA 1      2.17    2.61   0.44  20.5   0.84    0.37  17.1  increase   100
#  4 NGS   FA 1     40.5    42.8    2.31   5.7   0.65    1.51   3.73 increase   100
#  5 GMS   FA 1      1.62    1.94   0.32  19.7   0.81    0.26  16.0  increase   100
#  6 HW    FA 2     76.3    78.0    1.71   2.24  0.8     1.37   1.79 increase   100
#  7 HIS   FA 2     74.7    74.0   -0.66  -0.89  0.79   -0.52  -0.7  increase     0
#  8 PH    FA 3     86.5    86.4   -0.1   -0.11  0.78   -0.08  -0.09 decrease   100
#  9 SH    FA 3     77.8    77.0   -0.85  -1.09  0.82   -0.7   -0.9  decrease   100
# 10 FLH   FA 3     60.4    58.4   -2.03  -3.36  0.83   -1.68  -2.78 decrease   100
# 11 FLO   FA 4     60.4    59     -1.45  -2.41  0.92   -1.34  -2.22 decrease   100
# 12 DIS   FA 4      2.84    3.14   0.3   10.6   0.9     0.27   9.53 increase   100
# 13 NGSP  FA 4      2.62    2.66   0.05   1.74  0.52    0.02   0.9  increase   100
# 14 GY    FA 5   4380.   4715.   335.     7.65  0.72  241.     5.5  increase   100
```

##### 1.5.3 Coincidence index

We can use the function `coincidence_index()` to compute a coincidence index (Hamblin & Zimmermann, 1986) between multi-trait indexes.

```
coincidence_index(mgidi_index,
                  mgidi_fix,
                  mgidi_mean,
                  fai,
                  smith_index,
                  smith_index_colin,
                  total = 44)
# ------------------------------------------------------------------------------
# Coincidence index and common genotypes
# ------------------------------------------------------------------------------
# # A tibble: 15 x 5
#    V1          V2                  index common genotypes
#    <chr>       <chr>               <dbl>  <int> <chr>
#  1 mgidi_index mgidi_fix         100          7 G12,G37,G18,G8,G11,G9,G32
#  2 mgidi_index mgidi_mean        100          7 G12,G37,G18,G8,G11,G9,G32
#  3 mgidi_index fai                48.98       4 G12,G37,G18,G32
#  4 mgidi_index smith_index        -2.041      1 G32
```

```
#  5 mgidi_index smith_index_colin  31.97       3 G18,G11,G32
#  6 mgidi_fix   mgidi_mean        100          7 G12,G37,G18,G8,G11,G9,G32
#  7 mgidi_fix   fai                48.98       4 G12,G37,G18,G32
#  8 mgidi_fix   smith_index        -2.041      1 G32
#  9 mgidi_fix   smith_index_colin  31.97       3 G18,G11,G32
# 10 mgidi_mean  fai                48.98       4 G12,G37,G18,G32
# 11 mgidi_mean  smith_index        -2.041      1 G32
# 12 mgidi_mean  smith_index_colin  31.97       3 G18,G11,G32
# 13 fai         smith_index        14.97       2 G32,G7
# 14 fai         smith_index_colin  31.97       3 G18,G32,G7
# 15 smith_index smith_index_colin  31.97       3 G3,G7,G32
```

### 2 Appendix B

#### 2.1 Contribution of factor to the MGIDI of all genotypes

```
plot(mgidi_index,
     type = "contribution",
     genotypes = "all",
     n.dodge = 2,
     width.bar = 1,
     size.line = 0.1)
```

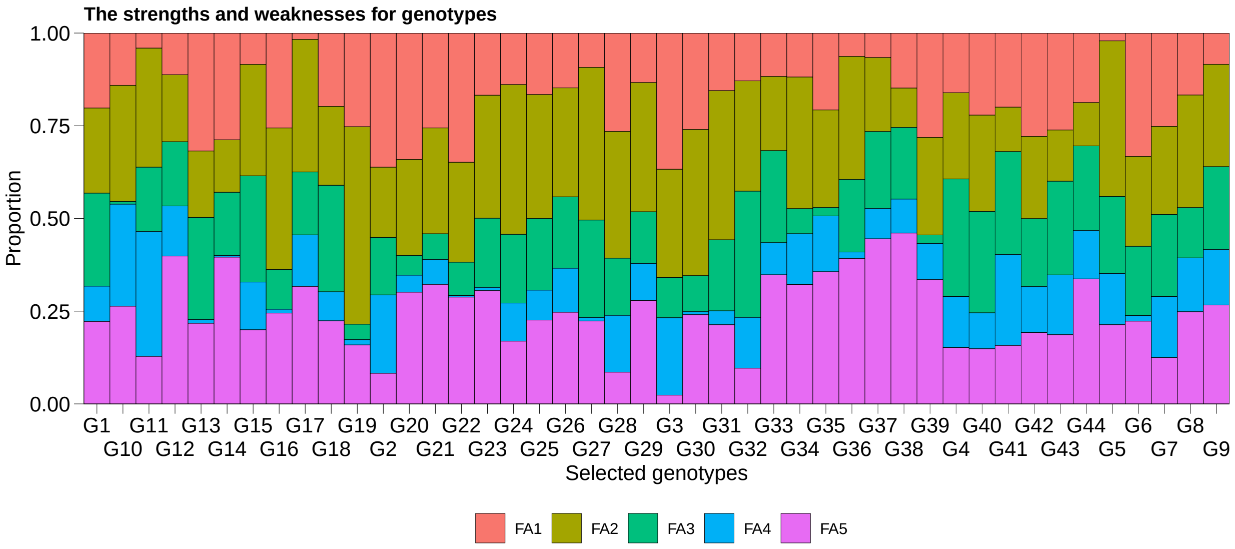

Figure S 2: Contribution of the factors to the MGIDI index of all studied genotypes.

#### 2.2 Phenotypic means for variables to decrease

```
# Create a helper function
plot_selected <- function(data,
                          gen,
                          var,
                          xlab = "Genotype",
                          ylab = expression(paste("Grain yield (kg ha"^-1,")"))){
  ovmean <- desc_stat(data, {{var}}, stats = "mean") %>% pull()
  plot_bars(data, {{gen}}, {{var}},
            xlab = xlab,
            ylab = ylab,
            size.text = 8,
            size.line = 0.2,
            width.bar = 1,
            fill.bar = c(rep("gray", 2),
                         rep("darkgoldenrod", 2),
                         rep("gray", 5),
                         "darkgoldenrod",
                         rep("gray", 15),
                         "darkgoldenrod",
                         rep("gray", 4),
                         "darkgoldenrod",
                         rep("gray", 11),
                         rep("darkgoldenrod", 2)),
            n.dodge = 2,
            order = "asce") +
    geom_hline(yintercept = ovmean, linetype = 2, color = "red", size = 0.2)
}
```

```
plot_selected(data, GEN, FLO, ylab = "Days to flowering")
```

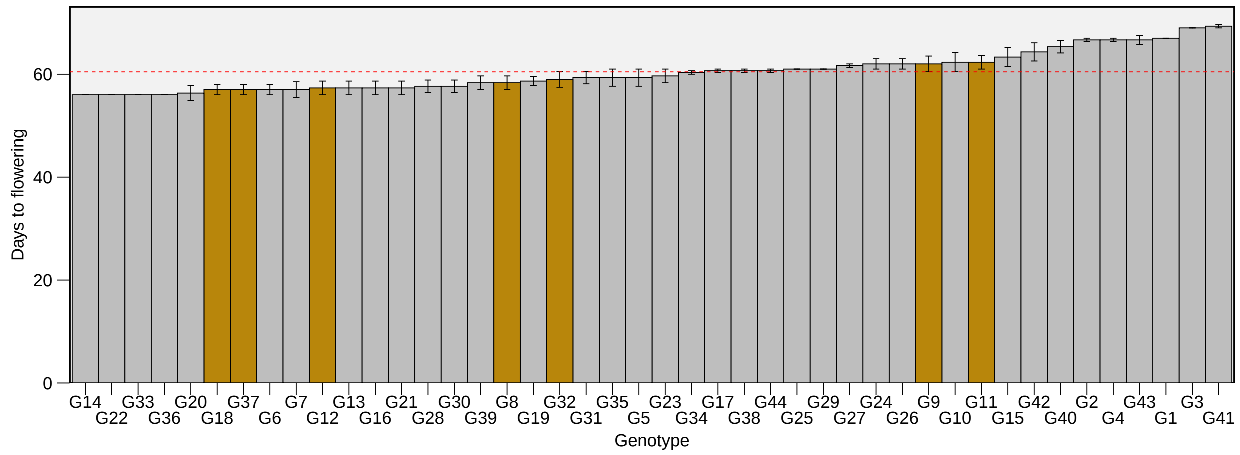

Figure S 3: Days to flowering for the evaluated genotypes. The selected genotypes by the MGIDI index are highlighted in dark golden color. Bars show the mean ± SE. N = 3.

```r
plot_selected(data, GEN, PH, ylab = "Plant heigth (cm)")
```

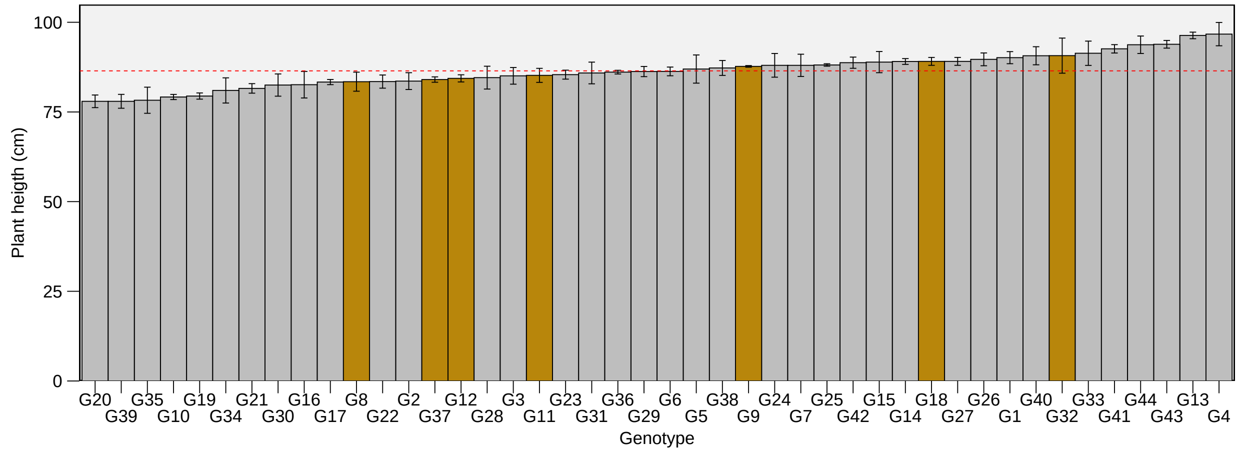

Figure S 4: Plant heigth for the evaluated genotypes. The selected genotypes by the MGIDI index are highlighted in dark golden color. Bars show the mean ± SE. N = 3.

```r
plot_selected(data, GEN, SH, ylab = "Spike height (cm)")
```

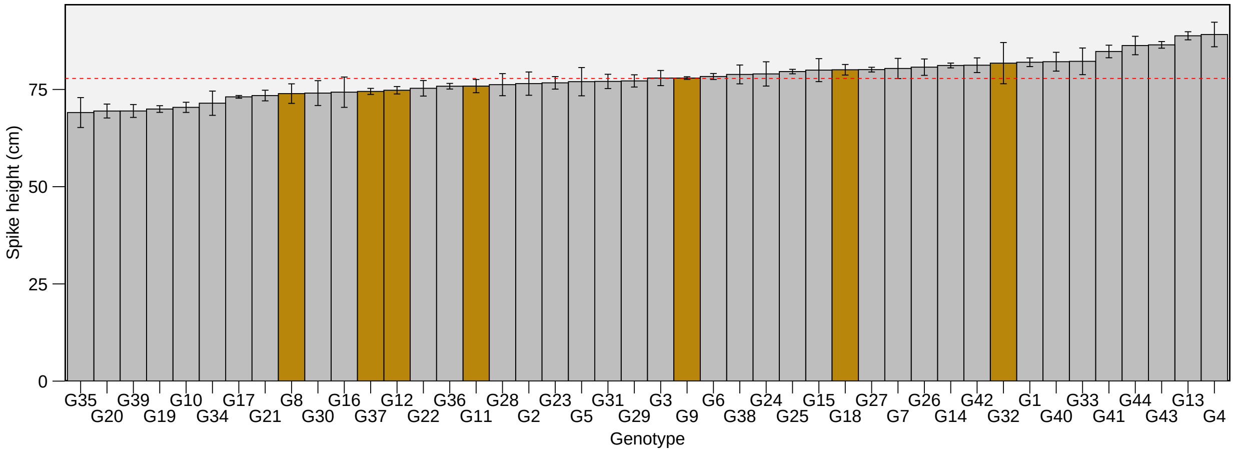

Figure S 5: Spike height for the evaluated genotypes. The selected genotypes by the MGIDI index are highlighted in dark golden color. Bars show the mean ± SE. N = 3.

```r
plot_selected(data, GEN, FLH, ylab = "Flag leaf height (cm)")
```

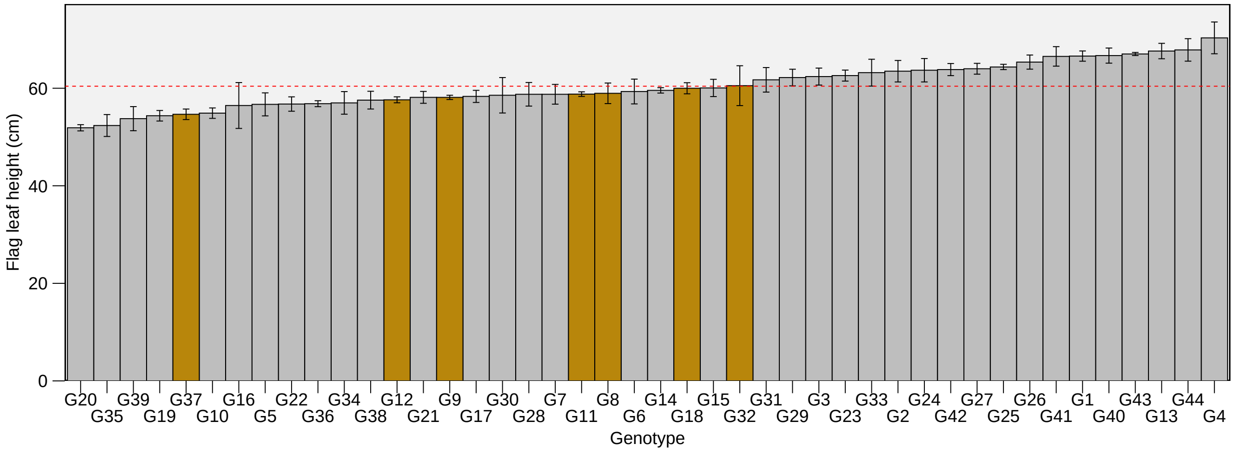

Figure S 6: Flag leaf height for the evaluated genotypes. The selected genotypes by the MGIDI index are highlighted in dark golden color. Bars show the mean ± SE. N = 3.

#### 2.3 Phenotypic means for variables to increase

```r
plot_selected(data, GEN, DIS, ylab = "Disease")
```

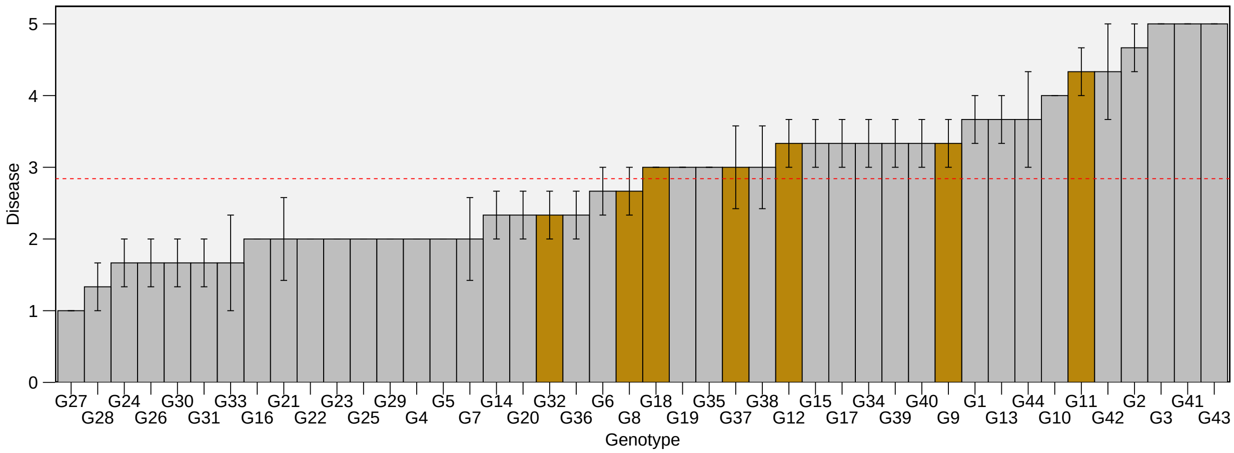

Figure S 7: Disease for the evaluated genotypes. The selected genotypes by the MGIDI index are highlighted in dark golden color. Bars show the mean ± SE. N = 3.

```r
plot_selected(data, GEN, HW, ylab = "Hectoliter weight")
```

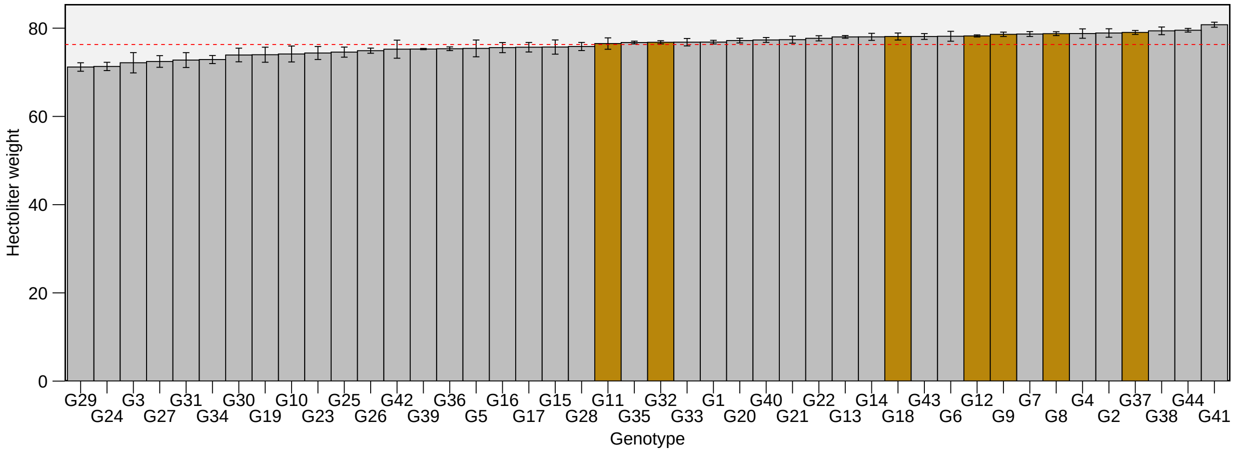

Figure S 8: Hectoliter weight for the evaluated genotypes. The selected genotypes by the MGIDI index are highlighted in dark golden color. Bars show the mean ± SE. N = 3.

```r
plot_selected(data, GEN, NSS, ylab = "Number of spiklets per spike")
```

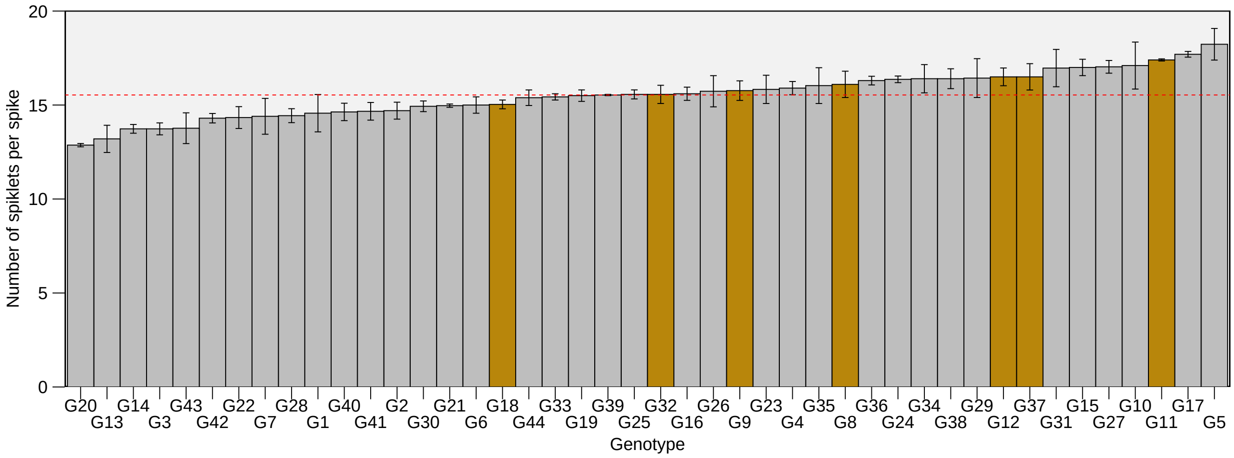

Figure S 9: Number of spiklets per spike for the evaluated genotypes. The selected genotypes by the MGIDI index are highlighted in dark golden color. Bars show the mean ± SE. N = 3.

```
plot_selected(data, GEN, NGSP, ylab = "Number of grains per spiklet")
```

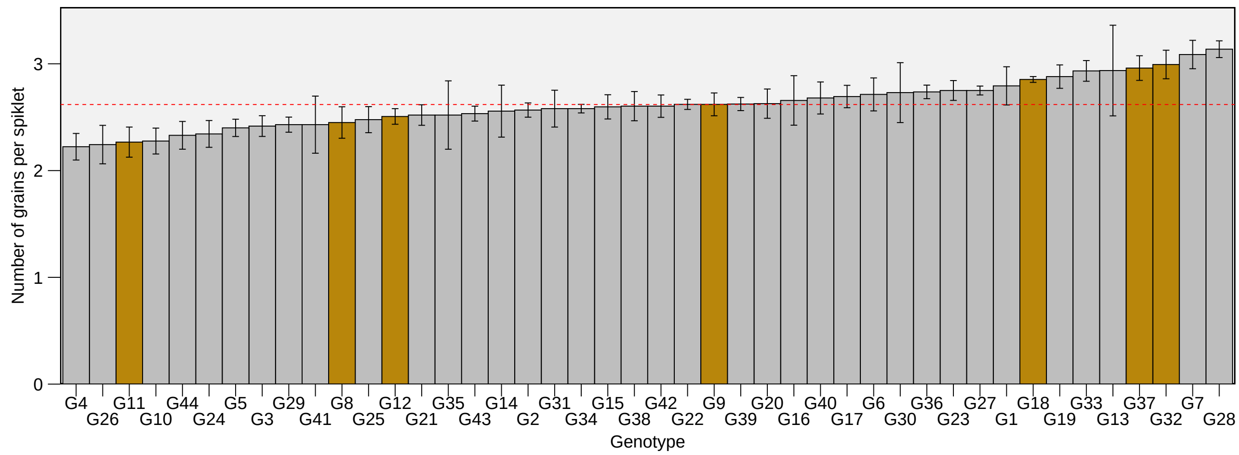

Figure S 10: Number of grains per spiklet for the evaluated genotypes. The selected genotypes by the MGIDI index are highlighted in dark golden color. Bars show the mean ± SE. N = 3.

```
plot_selected(data, GEN, SL, ylab = "Spike length")
```

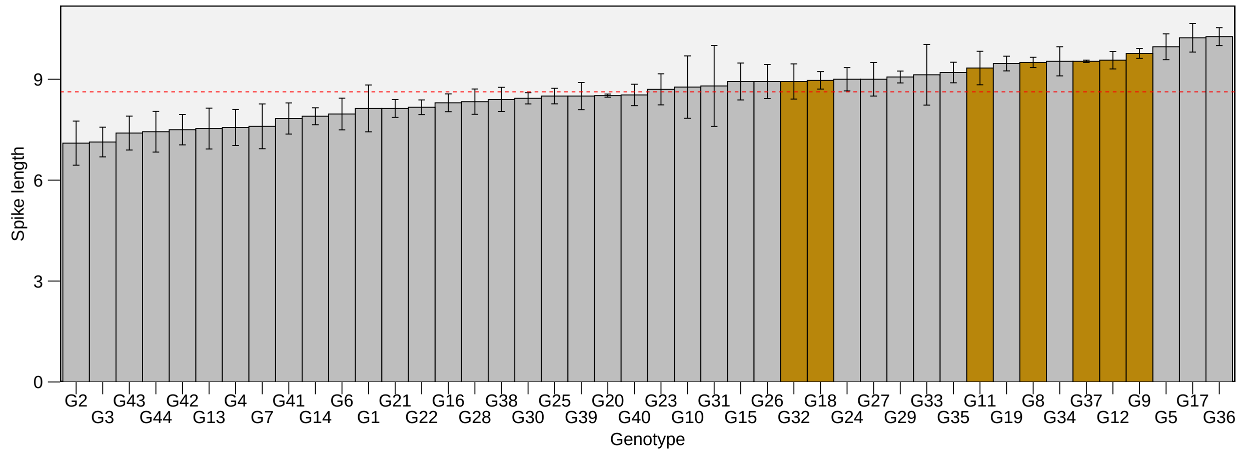

Figure S 11: Spike length for the evaluated genotypes. The selected genotypes by the MGIDI index are highlighted in dark golden color. Bars show the mean ± SE. N = 3.

```r
plot_selected(data, GEN, SW, ylab = "Spike weight")
```

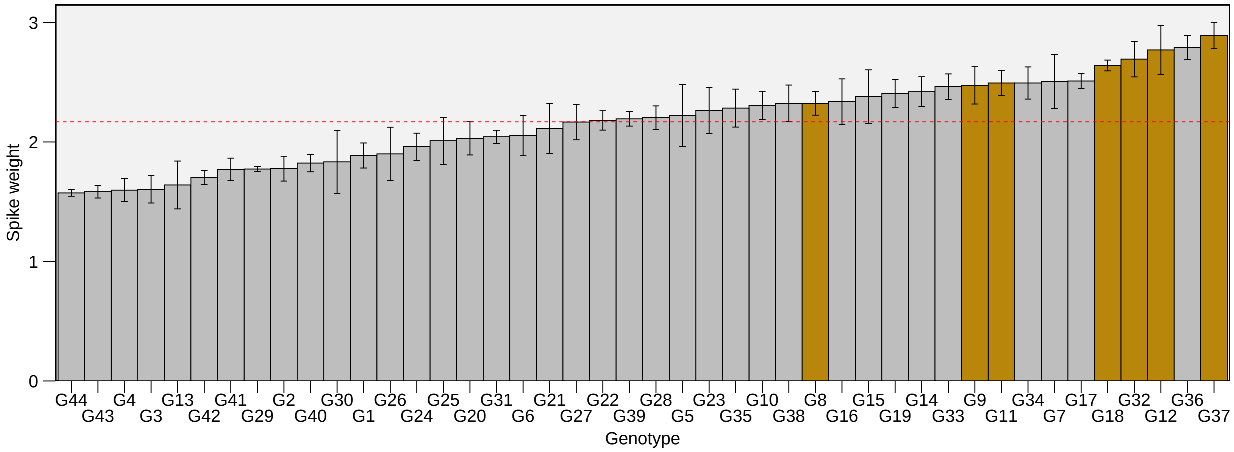

Figure S 12: Spike weight for the evaluated genotypes. The selected genotypes by the MGIDI index are highlighted in dark golden color. Bars show the mean ± SE. N = 3.

```r
plot_selected(data, GEN, NGS, ylab = "Number of grains per spike")
```

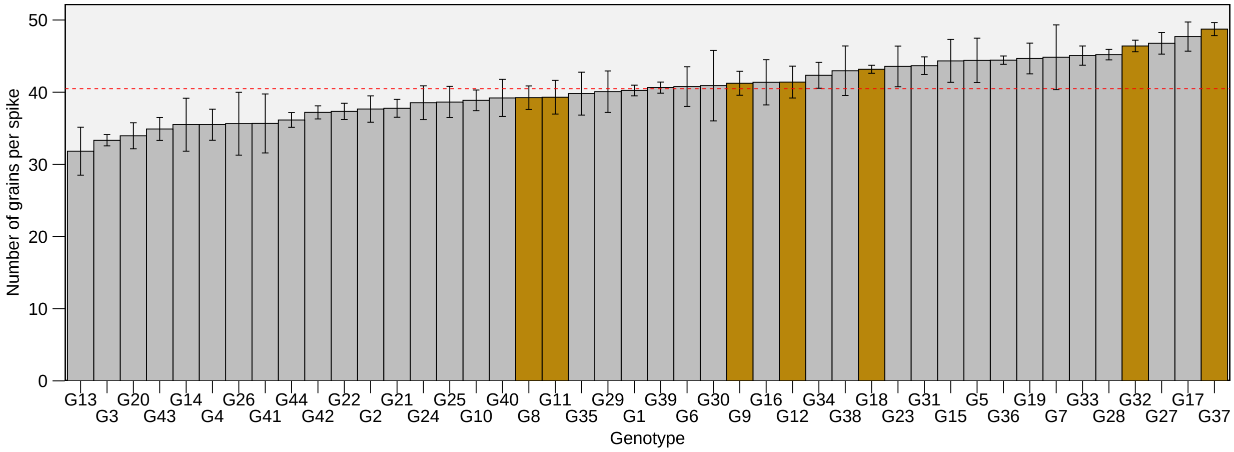

Figure S 13: Number of grains per spike for the evaluated genotypes. The selected genotypes by the MGIDI index are highlighted in dark golden color. Bars show the mean ± SE. N = 3.

```
plot_selected(data, GEN, GMS, ylab = "Grain mass per spike")
```

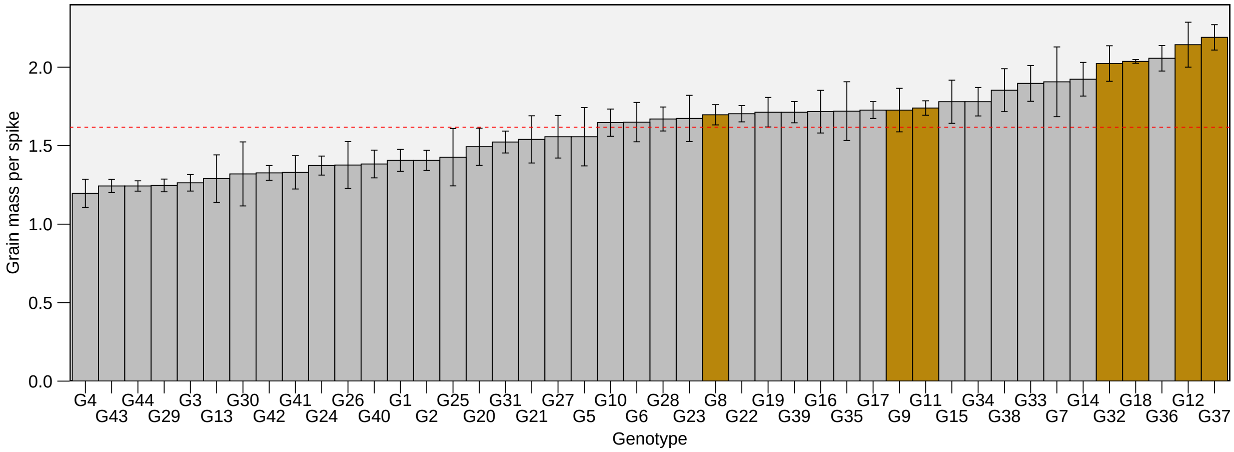

Figure S 14: Grain mass per spike for the evaluated genotypes. The selected genotypes by the MGIDI index are highlighted in dark golden color. Bars show the mean ± SE. N = 3.

```
plot_selected(data, GEN, HIS, ylab = "Harvest index of the spike")
```

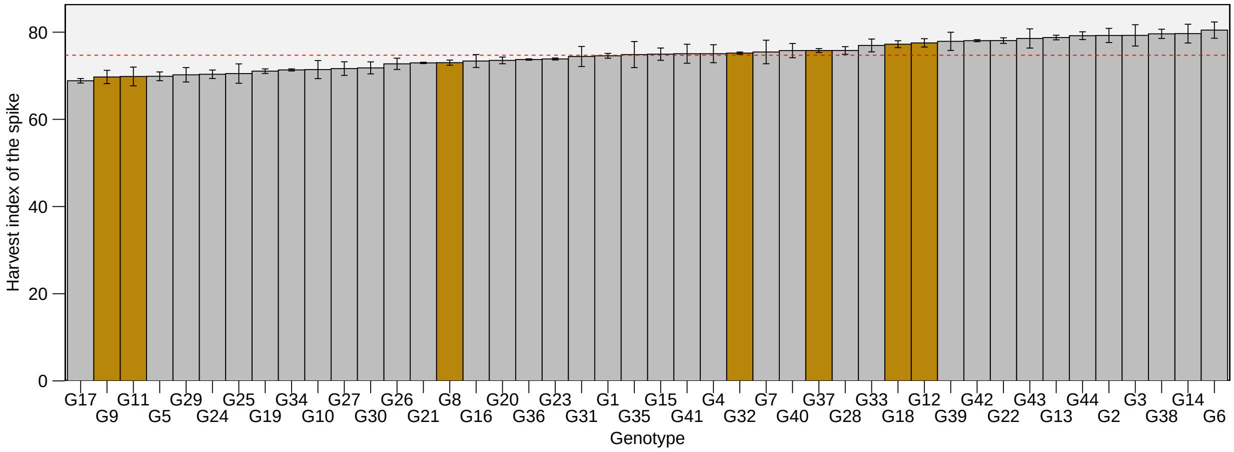

Figure S 15: Harvest index of the spike for the evaluated genotypes. The selected genotypes by the MGIDI index are highlighted in dark golden color. Bars show the mean ± SE. N = 3.

#### 2.4 Genotype ranking for the FAI-BLUP and Smith-Hazel indexes.

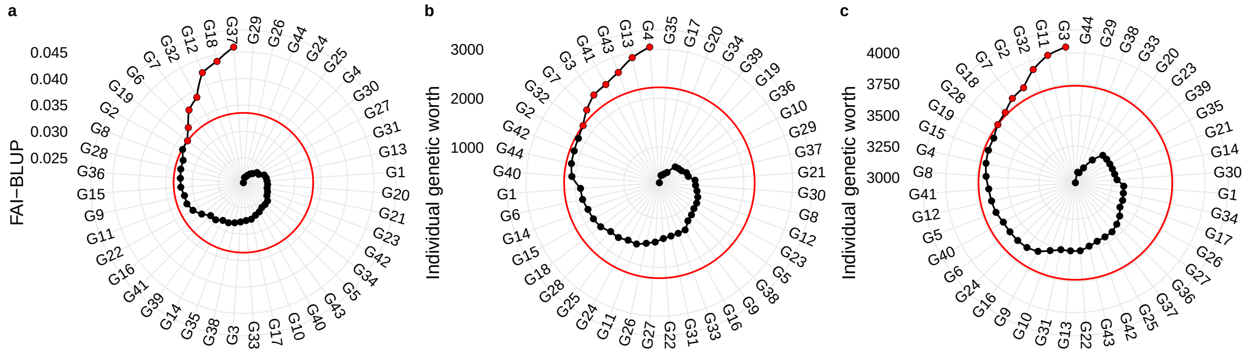

Figure S 16: Genotype ranking for the FAI-BLUP (a), Smith-Hazel with all traits (b), and after removing the traits (c).

### 3 Appendix C

Table S 1: Code, description and goal for selection of the traits evaluated in 44 wheat genotypes

| Code | Description | Unit | Goal |
|---|---|---|---|
| DF | Days to flowering | days | decrease |
| PH | Plant height | cm | decrease |
| SH | Spike height | cm | decrease |
| FLH | Flag leaf height | cm | decrease |
| DIS | Disease note | 0-5 | increase |
| GY | Grain yield | kg $ha^{-1}$ | increase |
| HW | hectoliter weight | kg $hL^{-1}$ | increase |
| NSS | number of spikelets per spike | count | increase |
| NGS | number of grains per spikelets | count | increase |
| SW | spike weight | g | increase |
| NGE | number of grains per spike | count | increase |
| GMS | grain mass per spike | g | increase |
| HIE | harvest index of the spike | GMS/SW | increase |

Table S 2: Deviance analysis, estimated variance components and genetic parameters for 14 agronomic traits evaluated in 44 wheat genotypes

| Traits[a] | $\hat{\sigma}_g^2$(%) | $h^2$ | $CV_g$ | $CV_r$ | p-value |
|---|---|---|---|---|---|
| DIS | 0.98(74.2) | 0.90 | 34.84 | 20.53 | 1.36e-18 |
| FLH | 16.65(61.3) | 0.83 | 6.75 | 5.36 | 6.20e-12 |
| GMS | 0.06(59) | 0.81 | 14.74 | 12.30 | 5.18e-11 |
| GY | 157688(46.1) | 0.72 | 9.07 | 9.81 | 5.71e-07 |
| HIS | 8.47(55.3) | 0.79 | 3.90 | 3.50 | 1.08e-09 |
| HW | 4.47(57.3) | 0.80 | 2.77 | 2.39 | 2.19e-10 |
| NGS | 11.05(38.8) | 0.65 | 8.21 | 10.32 | 2.89e-05 |
| NGSP | 0.03(26.1) | 0.51 | 6.10 | 10.26 | 4.83e-03 |
| NSS | 1.12(52.1) | 0.77 | 6.80 | 6.53 | 1.15e-08 |
| PH | 16.89(54.7) | 0.78 | 4.75 | 4.32 | 1.66e-09 |
| SH | 20.44(60.4) | 0.82 | 5.81 | 4.71 | 1.47e-11 |
| SL | 0.45(40.2) | 0.67 | 7.80 | 9.52 | 1.45e-05 |
| SW | 0.11(63) | 0.84 | 15.07 | 11.56 | 1.29e-12 |

[a] See Table S1 for the full trait names.

Table S 3: Eigenvalues, explained variance, factorial loadings after varimax rotation, and communalities obtained in the factor analysis

| Trait[a] | FA1 | FA2 | FA3 | FA4 | FA5 |
|---|---|---|---|---|---|
| NSS | **-0.876**[b] | -0.260 | 0.055 | -0.198 | 0.068 |
| SL | **-0.843** | -0.204 | 0.294 | 0.118 | 0.115 |
| SW | **-0.709** | 0.257 | 0.392 | 0.473 | -0.028 |
| NGS | **-0.716** | -0.096 | 0.091 | 0.539 | -0.086 |
| GMS | **-0.578** | 0.426 | 0.357 | 0.542 | -0.010 |
| HW | 0.079 | **0.847** | -0.177 | 0.051 | 0.040 |
| HIS | 0.541 | **0.653** | -0.156 | 0.170 | 0.015 |
| PH | -0.075 | -0.171 | **0.966** | 0.013 | -0.008 |
| SH | -0.209 | -0.192 | **0.947** | 0.032 | 0.011 |
| FLH | -0.265 | 0.066 | **0.900** | 0.243 | 0.030 |
| FLO | -0.124 | 0.013 | 0.400 | **0.724** | 0.355 |
| DIS | 0.142 | 0.563 | 0.034 | **-0.617** | -0.294 |
| NGSP | 0.033 | 0.143 | -0.007 | **0.867** | -0.167 |
| GY | 0.033 | -0.003 | -0.005 | -0.006 | **-0.933** |
| Eigenvalues[c] | 5.570 | 2.410 | 1.820 | 1.380 | 1.010 |
| Variance | 39.760 | 17.230 | 12.990 | 9.840 | 7.200 |
| Accumulated | 39.760 | 56.990 | 69.980 | 79.820 | 87.020 |

[a] See Table S1 for the full trait names.
[b] Bold values indicate the variables grouped within each factor.
[c] See the entirely list of eigenvalues as Appendix A1.4.1.5.

Table S 4: Coincidence index (CI) of genotype selection for each pair of indexes evaluated

| Index 1 | Index 2 | CI | Shared genotypes |
|---|---|---|---|
| MGIDI | FAI-BLUP | 48.98 | G12,G37,G18,G32 |
| MGIDI | SH-1 | -2.05 | G32 |
| MGIDI | SH-2 | 31.98 | G18,G11,G32 |
| FAI-BLUP | SH-1 | 14.96 | G32,G7 |
| FAI-BLUP | SH-2 | 48.97 | G18,G32,G7,G2 |
| SH-1 | SH-2 | 31.97 | G3,G7,G32 |

### References

Bengtsson, H. (2020). *Future.apply: Apply function to elements in parallel using futures*. Retrieved from https://CRAN.R-project.org/package=future.apply

Hamblin, J., & Zimmermann, M. J. de O. (1986). Breeding Common Bean for Yield in Mixtures. In *Plant breeding reviews* (pp. 245–272). Hoboken, NJ, USA: John Wiley & Sons, Inc. doi:10.1002/9781118061015.ch8

Olivoto, T., Carvalho, I. R., Nardino, M., Ferrari, M., Pelegrin, A. J. de, Follmann, D. N., … Souza, V. Q. de. (2016). Sulfur and nitrogen effects on industrial quality and grain yield of wheat. *Revista de Ciências Agroveterinárias*, *15*(1), 24–33. doi:10.5965/223811711512016024

Olivoto, T., & Lúcio, A. D. (2020). metan: an R package for multi-environment trial analysis. *Methods in Ecology and Evolution*, *11*(6), 783–789. doi:10.1111/2041-210X.13384

Rocha, J. R. do A. S. de C., Machado, J. C., & Carneiro, P. C. S. (2018). Multitrait index based on factor analysis and ideotype-design: proposal and application on elephant grass breeding for bioenergy. *GCB Bioenergy*, *10*(1), 52–60. doi:10.1111/gcbb.12443

Smiderle, É., Furtini, I., Botelho, F. B., Resende, M. P., Botelho, R. T., Colombari Filho, J. M., … Utumi, M. (2019). Index selection for multiple traits in upland rice progenies. *Revista de Ciências Agrárias*, *42*(1), 1–10. doi:10.19084/RCA18059

Wickham, H., Averick, M., Bryan, J., Chang, W., McGowan, L., François, R., … Yutani, H. (2019). Welcome to the Tidyverse. *Journal of Open Source Software*, *4*(43), 1686. doi:10.21105/joss.01686
